# STEP-PTMs: Sequential TMT-based Enrichment and Profiling of Post-Translational Modifications

**DOI:** 10.64898/2026.08.18.745386

**Authors:** Lucrezia Criscuolo, Sofie B. Elmkvist, Arkadiusz Nawrocki, Lene A. Jakobsen, Pia Jensen, Peter T. Jensen, Honggang Huang, Jesper F. Havelund, Nils J. Færgeman, Giuseppe Palmisano, Helle Bogetofte, Martin R. Larsen

## Abstract

Comprehensive characterization of protein abundance and multiple post-translational modifications (PTMs) from the same biological samples is essential for understanding cellular regulation and PTM crosstalk but remains analytically challenging. Here, we present STEP-PTM (Sequential Tag-based Enrichment of Post-Translational Modifications), a modular TMT-multiplexed workflow that enables integrated quantitative analysis of the proteome, metabolome and multiple PTM classes from a single peptide preparation. Proteins are digested, isobarically labeled using tandem mass tags (TMT), and combined into a single multiplexed peptide pool prior to sequential PTM enrichment, thereby minimizing technical variability, reducing sample requirements and facilitating direct quantitative integration across datasets.

STEP-PTM supports flexible sequential enrichment of phosphopeptides, peptides containing free and reversibly modified cysteines, sialylated N-linked glycopeptides, lysine-acetylated peptides and S-palmitoylated peptides, while preserving non-modified peptides for global proteome analysis. PTM-specific database searches further improve identification confidence and quantitative accuracy, and the modular workflow can readily be adapted by incorporating or omitting enrichment modules according to the biological question.

Application of STEP-PTM to TMT16-plex cerebral brain organoids enabled the quantification of 10,413 proteins, 2,969 metabolites, 19,655 phosphopeptides, 28,876 peptides containing reversibly modified cysteines, 9,723 peptides containing free cysteines, 1,716 intact sialylated N-linked glycopeptides and 771 lysine-acetylated peptides from the same biological samples. We further demonstrate the applicability of the workflow to multiple mouse tissues, highlighting its broad utility for integrated systems-level characterization of protein expression and PTM regulation across diverse biological models.

## Introduction

Today, it is well established that analysis of protein abundance alone provides only a partial view of cellular regulation, as proteome complexity is greatly expanded by reversible and irreversible post-translational modifications (PTMs) [1]. Together with changes in protein abundance, PTMs regulate protein conformation, activity, stability, localization and interactions with DNA, RNA and other proteins, thereby controlling virtually every aspect of cellular physiology [2–5].

Public PTM databases now contain more than 270,000 experimentally identified PTM sites in human proteins, with this number continuing to grow. To date, hundreds of chemically distinct PTMs have been described, highlighting the extraordinary complexity of proteome regulation [6, 7]. The human proteome contains numerous enzyme families dedicated to the installation, removal and processing of PTMs, including kinases, phosphatases, glycosyltransferases, acetyltransferases, deacetylases, ubiquitin ligases, deubiquitinases and proteases. [8, 9]. In addition, chemically reactive amino acids such as cysteine undergo numerous reversible oxidative modifications that play fundamental roles in regulating protein function and enzymatic activity [10–15].

The functional complexity of the PTMome extends beyond the large number of modification types. Many PTMs are highly dynamic and reversible, enabling rapid spatiotemporal regulation of signaling pathways, cellular differentiation, apoptosis and numerous other biological processes [16, 17]. Furthermore, multiple PTMs frequently coexist on the same protein or within the same signalling pathway, resulting in PTM crosstalk where one modification influences the occurrence or function of another [6, 18–20]. Consequently, comprehensive characterization of multiple PTMs has become increasingly important for understanding cellular regulation in both physiology and disease. Aberrant PTM patterns have been associated with numerous pathological conditions, including cancer, neurodegenerative disorders and metabolic diseases, making PTMs attractive sources of disease biomarkers and therapeutic targets [5, 17, 21].

Mass spectrometry has become the gold standard for large-scale proteomics and PTM analysis [3, 22]. However, comprehensive characterization of multiple PTMs remains analytically challenging because most PTMs occur at low stoichiometry, exhibit diverse physicochemical properties and often generate complex MS/MS spectra. In addition, PTM analysis is influenced by digestion efficiency, peptide ionization, chromatographic behavior, PTM stability and the computational challenges associated with database searching [23, 24]. Consequently, selective enrichment of modified peptides before LC-MS/MS has become essential for comprehensive PTM analysis.

Numerous enrichment strategies have therefore been developed for individual PTMs. Immunoaffinity-based approaches use modification-specific antibodies to isolate peptides containing selected PTMs, such as lysine acetylation or ubiquitination [25–27], whereas affinity chromatography exploits the physicochemical properties of modifications including phosphorylation and sialylated N-glycosylation using IMAC or TiO₂ chromatography [28–34]. Other strategies rely on chemical derivatization of specific PTMs before enrichment [25, 35–43]. One example is the Cysteine-specific Phosphonate Adaptable Tag (CysPAT) [38], which enables selective labeling of free or reversibly modified cysteines for subsequent enrichment by TiO₂ or IMAC chromatography [38, 44]. While these methods have enabled deep characterization of individual PTMs, most workflows still focus on a single modification class together with the proteome.

The growing appreciation of PTM crosstalk has stimulated the development of workflows capable of profiling multiple PTMs within the same biological samples [20, 45]. Our laboratory previously developed a sequential TiO₂-based strategy for simultaneous analysis of phosphorylation, lysine acetylation and sialylated N-linked glycosylation [46], later extended with SIMAC to separate mono- and multiphosphorylated peptides [47]. Carr and colleagues subsequently introduced SEPTM, combining phosphorylation, ubiquitination and lysine acetylation using SILAC-based quantification

[48]), followed by a multiplexed TMT workflow for integrated proteome and phosphoproteome analysis [49]. More recently, we developed TiCPG, enabling quantitative analysis of phosphopeptides, sialylated N-glycopeptides and reversibly modified cysteines using iTRAQ labeling from as little as 100 μg protein per sample [50]. Recently, Xu et al. introduced the MoSAIC workflow, representing an important advance toward integrated analysis of multiple PTMs by combining phosphorylation, glycosylation, acetylation and ubiquitination within a modular experimental framework [51]. In this strategy, however, each PTM is enriched independently before isobaric labeling. Consequently, quantitative performance depends on the reproducibility of multiple independent enrichment procedures, labeling reactions, sample normalization steps and LC-MS/MS analyses.

Despite these advances, protocols enabling comprehensive large-scale analysis of multiple PTMs from the same biological samples remain limited. Such workflows are essential for advancing our understanding of cell signaling, PTM crosstalk and disease mechanisms by enabling direct quantitative integration of multiple regulatory layers within the same experiment.

To address this need, we developed STEP-PTM (Sequential Tag-based Enrichment of Post-Translational Modifications), a modular workflow in which proteins are digested, isobarically labeled and combined into a single multiplexed peptide pool before PTM enrichment (**Figure 1**). All subsequent enrichment, fractionation and LC-MS/MS analyses are therefore performed on the pooled sample, minimizing technical variability introduced after multiplexing, reducing sample requirements and enabling direct quantitative integration of the proteome with multiple PTM datasets. Furthermore, PTM-specific database searches can be performed for each enriched fraction, improving identification confidence and quantitative accuracy. Owing to its modular design, STEP-PTM can readily be adapted by incorporating or omitting individual enrichment modules according to the biological question and sample type.

**Figure 1.**
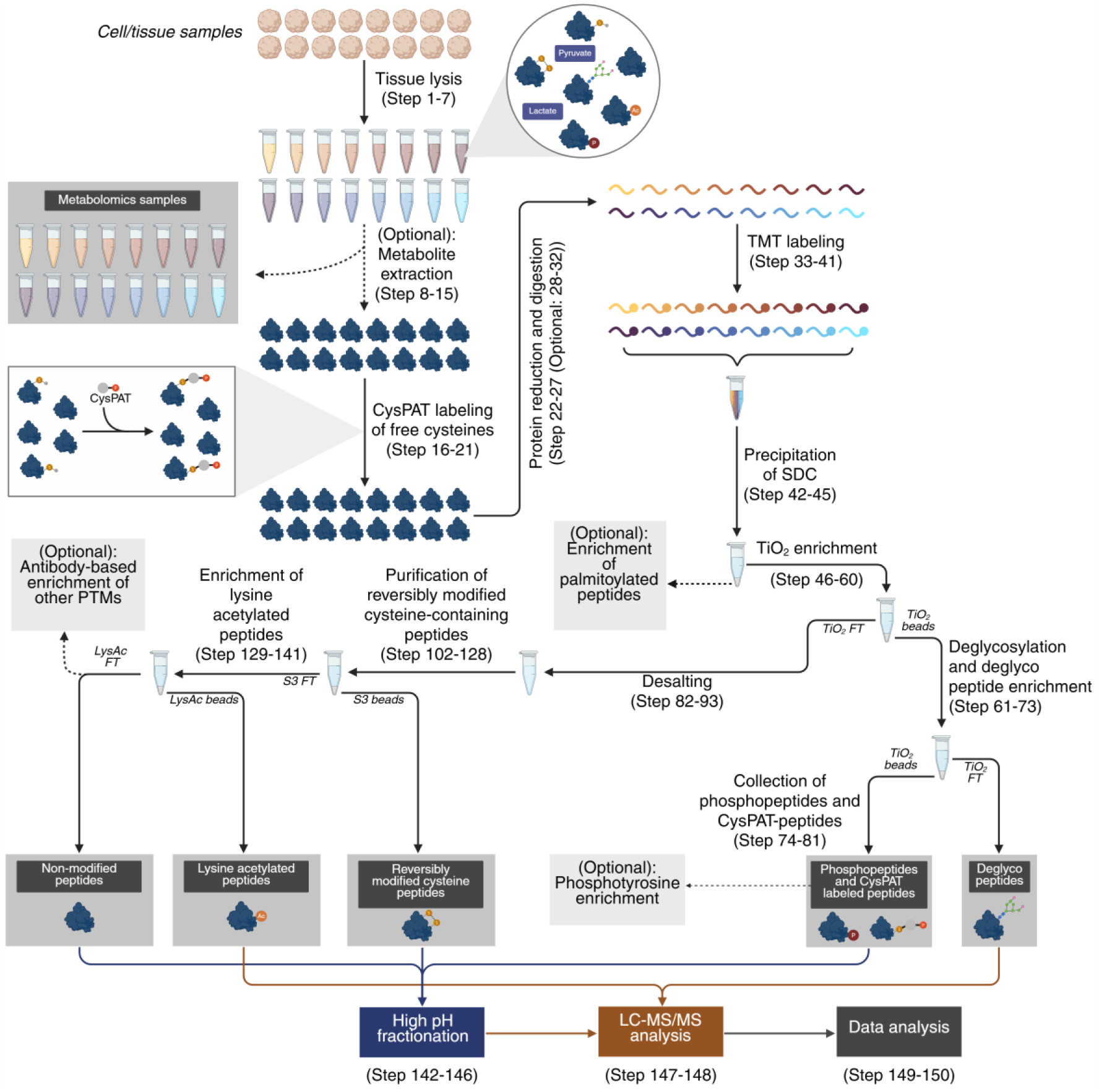
| Overview of the STEP-PTM sample preparation workflow for global multiplexed proteomic and PTMomic analysis of biological samples using isobaric labelling. The workflow includes optional and recommended steps for sample preparation, including protein extraction and digestion, followed by isobaric labelling using tandem mass tags (TMT) for relative quantification between different biological samples and sequential enrichment of different types of PTMs, of which one or more PTM enrichment steps can be included in an experiment according to the experimental design.

In its current implementation, STEP-PTM enables the comprehensive identification and quantification of phosphopeptides, peptides containing free and reversibly modified cysteines, lysine-acetylated peptides, and sialylated N-linked glycopeptides, together with global proteome and metabolome analyses from the same biological samples. The workflow can readily be expanded to include additional PTM modules, such as enrichment of S-palmitoylated peptides [52].

### Development of the protocol

For decades, the fundamental role of PTMs in both physiological and pathological mechanisms have been well-established. Our group, along with others, has developed various mass spectrometry-based approaches to study individual or a limited number of PTMs. However, devising a robust strategy for the large-scale identification and quantification of multiple PTMs simultaneously remains a significant challenge. In 2012, our group developed a protocol to simultaneously profile protein phosphorylation, lysine acetylation, and N-linked sialylated glycosylation [46]. Specifically, the method proposed involved the use of TiO_2_ to enrich phosphopeptides and sialylated N-linked glycopeptides. Then, after TiO_2_ chromatography, K(Ac) antibodies immunoprecipitated the lysine acetylated peptides contained in the first TiO_2_ flow-through. This method was later combined with SIMAC to separate mono-phosphorylated peptides from multiply phosphorylated peptides [47].

This method offered the simultaneous coverage of the proteome and three PTMs within a biological sample, but is lacking a multiplexing strategy, limiting its ability to compare multiple conditions or samples.

In 2013, Carr *et al.* proposed a method for serial enrichment of PTMs (SEPTM), that included the analysis of phosphorylation, ubiquitination, and lysine acetylation from the same biological sample [48]. The method is based on the use of affinity-based techniques, such as IMAC for phosphopeptide enrichment, K(GG) antibody beads for enrichment of ubiquitinylated peptides, and K(Ac) antibody beads for enrichment of lysine-acetylated peptides. To gain reliable and robust quantification of PTMs, SEPTM involves the labelling of proteins with SILAC (Stable Isotope Labeling by Amino Acids in Cell Culture) [53, 54]. This approach produced good results in terms of peptide identifications, and the coverage of PTMs could be improved by adding a step of offline fractionation prior to the LC-MS/MS analysis. However, a significantly high starting material is required -approximately 1 mg of protein per SILAC label, increasing to 2.5 mg if offline fractionation is performed.

In 2018, Carr and colleagues published another protocol for multiplexed proteome and phosphoproteome analysis [49]. This method allowed a reproducible strategy for the analysis of tissues or cell lines, incorporating the use of TMT10 labeling for multiplexing and IMAC for phosphopeptide enrichment. The proteome and phosphoproteome coverages were further improved by offline fractionation, giving a high number of identifications per sample, and reliable quantifications. However, the starting material required was still considerable – 300 µg of peptide per sample, restricting the use for low abundant or precious samples.

In 2016, our group synthesized the Cysteine-specific Phosphonate Adaptable Tag (CysPAT) and developed a method to selectively label cysteine-containing peptides (Cys peptides) with CysPAT and simultaneously enrich Cys peptides and phosphopeptides using titanium dioxide (TiO_2_) chromatography and subsequently characterized their changes after 5 min epidermal growth factor (EGF) stimulation of HeLa cells [38]. Later in 2023, our group developed another mass spectrometry-based approach called TiCPG that enabled the enrichment of three PTMs, including phosphopeptides, sialylated N-glycopeptides and peptides with reversibly modified cysteines (RmCys) [50]. The strategy used CysPAT labeling of RmCys peptides, a TiO_2_ enrichment step to fractionate the non-modified peptides [38], the enriched fraction was followed by the deglycosylation of sialylated N-glycopeptides, which were then separated through a second TiO_2_ enrichment. The method, combining iTRAQ (Isobaric Tags for Relative and Absolute Quantitation) labeling for quantitation and LC-MS/MS, was suitable for quantitative characterization of three PTMs from a low amount of starting material -100 μg per iTRAQ channel.

Recently, Xu et al. introduced the MoSAIC workflow, representing an important advance toward integrated analysis of multiple PTMs by combining phosphorylation, glycosylation, acetylation and ubiquitination within a modular experimental framework [51]. In this workflow, each PTM is enriched independently from individual biological samples prior to isobaric labeling. Consequently, quantitative performance depends on the reproducibility and selectivity of multiple independent enrichment procedures, TMT labeling reactions, sample normalization steps and LC-MS/MS analyses. Because each PTM dataset is generated independently, quantitative integration across PTM classes requires comparison of multiple separately generated TMT experiments. Furthermore, individual PTM enrichments require separate sample aliquots, increasing the amount of starting material required for comprehensive PTM characterization.

Although this strategy provides broad PTM coverage, performing enrichment prior to sample multiplexing means that any variation introduced during enrichment directly contributes to quantitative variability between biological samples. Since affinity-based enrichment methods are never completely identical between independent experiments, highly reproducible enrichment procedures and increased biological replication become important for robust quantitative analysis.

To overcome these limitations, we developed STEP-PTM (Sequential Tag-based Enrichment of Post-Translational Modifications), a highly flexible workflow that introduces isobaric labeling immediately after protein digestion, allowing all biological samples to be combined into a single multiplexed peptide pool before any PTM enrichment is performed (**Figure 1**). Consequently, all subsequent enrichment procedures, washing steps, fractionation and LC-MS/MS analyses are performed only once on the pooled sample, ensuring identical experimental handling of every biological sample after multiplexing. This design minimizes technical variation introduced during PTM enrichment, simplifies normalization across experimental conditions and substantially reduces sample handling.

Another important advantage of STEP-PTM is the reduced requirement for starting material. Instead of dividing each biological sample into multiple aliquots for individual PTM enrichments, sequential enrichment of multiple PTM classes is performed from the same multiplexed peptide pool, enabling comprehensive characterization of the proteome together with phosphopeptides, reversibly modified and free cysteine-containing peptides, sialylated N-linked glycopeptides, S-palmitoylated peptides and lysine-acetylated peptides from as little as 30–100 μg of protein per sample, while supporting multiplexing of up to 18–32 (and potentially 35) biological samples within a single experiment with 100% overlap. In addition, the workflow can readily be expanded to include other PTMs using commercially available immunoaffinity reagents (e.g., PTMScan®, Cell Signaling Technology).

Because all PTM datasets originate from the same multiplexed peptide population, quantitative integration between the proteome and individual PTM datasets is straightforward and does not require comparison of independently normalized TMT experiments. Furthermore, each enriched fraction contains only one or a limited number of PTM classes, allowing database searches to be tailored specifically to the chemistry of each fraction. This minimizes unnecessary expansion of the search space while improving confidence in peptide identification and PTM site localization and secure a more reliable calculation of false discovery rates using validated programs such as Percolator (REF).

Finally, the protocol is designed as a modular workflow in which individual enrichment steps can be readily incorporated or omitted according to the biological question, providing a flexible platform for comprehensive PTM characterization across diverse biological samples, including cultured cells, organoids, complex tissues and liquid biopsies.

### Applications of the method

The strategy described here (STEP-PTM) has been developed to accommodate a wide range of applications, making it highly versatile for various research purposes and heterogeneous sample types. As reported in the *“Anticipated results*” section, this method was applicable to various biological samples, including 2D and 3D cell cultures, biofluids and complex tissues, such as cardiac tissue and liver [45, 50, 55].

Starting with a minimum of 50-100 µg of proteins per sample using TMT multiplexing, this method allows for reproducible and simultaneous identification and quantification of multiple PTMs on a global scale within a single experiment. Using higher multiplexing lower amount of starting material (e.g., 30 µg) is possible. Additionally, the protocol enables a comprehensive profiling of the proteome and the opportunity to also include analysis of the metabolome. Furthermore, if specific sub-proteomes, such as membrane-associated proteins or organelle proteins is a research focus, then the STEP-PTM method can be applied after purification of these sub-proteomes.

The high flexibility of our protocol allows researchers to tailor the protocol, either including or omitting enrichment steps for specific PTMs, based on experimental needs.

Additionally, we displayed the necessity of profiling multiple PTMs in diverse contexts for the study of both biological functions and pathological mechanisms. Indeed, the use of TMT multiplexing supports the analysis of up to 18-32 samples in parallel, allowing for the simultaneous study of multiple conditions and biological replicates with no missing values.

Finally, the multiplexing approach also improves the sensitivity for detecting low abundant peptides, by mixing TMT multiplexed peptides and off-line fractionating peptides with similar physicochemical properties before LC-MS/MS analysis.

### Experimental design

Here, we present an optimized strategy to profile and quantify the proteome, multiple PTMs and the metabolome of complex biological samples such as neural organoids. The protocol was designed as individual steps, where each one can be easily integrated or excluded according to the user’s interest. Thus, the experiment setup can be changed and adjusted, considering the following points:

1. The number of samples and/or conditions per experiment is flexible. However, it is advisable to have a minimum of 1 mg of protein in total as starting material (if applying a TMT16 plex, this corresponds to 62.5 µg protein per sample). If the amount of starting material is lower than 0.8-1 mg, a decrease in coverage of the individual PTM peptides should be expected.
2. The protocol is described using TMT multiplexing, but other labeling strategies can be utilized (e.g. iTRAQ). Multiplexing can be omitted; however, the accuracy of the subsequent peptide quantitation would decrease, due to errors in the reproducibility of multiple enrichments and fractionation methods.
3. The metabolome analysis is optional, but if included, the users should be aware of some steps in the sample preparation workflow. It is crucial to avoid using PBS (Phosphate-buffered saline) in the sample preparation, instead employing ammonium acetate for both sample washing and collection (in cases involving cells or in vitro cultures). Also, the use of SDC (Sodium deoxycholate) will eliminate lipids from metabolomic analysis.
4. As reported in the protocol, as an optional step, it is possible to enrich S-palmitoylated peptides from the SDC pellet as described (“Precipitation of SDC”, steps 42-45) [52]. In this case, it is crucial to not quench the TMT reaction using hydroxylamine as this reagent will remove S-palmitoylation. Additionally, it is important to use 1-3 mM TCEP (Tris(2-carboxyethyl)phosphine hydrochloride solution) for reduction instead of DTT (Dithiothreitol) during sample preparation. An example of results obtained when including S-palmitoylated peptide enrichment is presented in the “*Anticipated results*” section.
5. After TMT labeling and mixing we are choosing a highly optimized TiO_2_ chromatographic method for enrichment of phosphopeptides, CysPAT peptides and sialylated N-glycopeptides, as the TiO_2_ resin is much more robust towards many low molecular contaminants in comparison to IMAC resin[56] and therefore an RP cleaning step is not necessary prior to enrichment.
6. After collection of the phosphopeptide and CysPAT labeled peptide fraction, it is possible to specifically separate tyrosine phosphorylated peptides from serine/threonine phosphorylated peptides using phosphotyrosine immunoaffinity enrichment with antibodies targeting pY [57, 58] or immobilized SH2 superbinder domains [59]. However, since phosphotyrosines are present in low stoichiometry, this step requires a larger amount of starting material for effective enrichment. We recommend having at least a total of 1.6 mg of peptide after combining TMT-labelled samples (corresponding to 100 μg peptide per TMT channel for a TMT 16-plex) as starting material prior to including the phosphotyrosine enrichment step.
7. It is possible to integrate the protocol with the study of additional PTMs, which can be enriched using antibody-based immunoaffinity enrichment. This includes ubiquitination, methylation, O-GlcNAcylation, succinylation and propylation, which can be enriched by the utilization of commercially available antibodies (e.g., PTMscan^®^ antibodies, Cell Signaling Technology).
8. To resolve the TMT-labeled peptides, the mass spectrometer used must be able to achieve a high resolution in the low mass range and relatively fast scanning. In this protocol, we used Orbitrap Eclipse Tribrid, Orbitrap Exploris 480 and Orbitrap Fusion Lumos instruments (Thermo Fisher Scientific). However, other MS instruments with the mentioned requirements for TMT multiplex analysis can be used.

## Materials

### REAGENTS

- Cells or tissues. (See the reagent setup for details)
  **!** CAUTION: All experiments with cell lines derived from humans as well as experiments with live animals should be performed according to institutional and national regulations.
- Sodium deoxycholate (SDC; Sigma-Aldrich, cat. no. D6750)
- Ammonium acetate (Sigma-Aldrich, cat. no. 73594)
- PhosSTOP (Phosphatase inhibitory cocktail tablets; Roche, cat. no. 04 906 837 001)
- Formic acid (FA; Merck, cat. no. 1.11670.0250) ▲ CRITICAL: Purchase LC- and MS-grade reagents
**!** CAUTION: FA is a health hazard compound, toxic and flammable. Avoid contact with skin and eyes, and avoid inhalation. Wear protective gloves and clothing. Use it in a chemical fume hood.
- Iodoacetic acid N-hydroxysuccinimide ester (SIA crosslinker; CovaChem, cat. no. 13408-1) ▴ CRITICAL: SIA crosslinker is unstable and light-sensitive. It should be kept in a cool and light protective place. SIA is also moisture-sensitive. To avoid moisture condensation onto the product, the vial must be equilibrated to room temperature before opening and sealed under argon.
**!** CAUTION: SIA crosslinker is toxic. Wear protective gloves and clothing. Use it in a chemical fume hood.
- 2-aminoethylphosphonic acid (2-AEP; Sigma-Aldrich, cat. no. 268674) ▲ CRITICAL: 2-AEP is unstable and light-sensitive, it should be kept in a cool and light protective place. To avoid moisture condensation onto the product, the vial must be equilibrated to room temperature before opening and sealed under argon.
- Dimethyl sulfoxide (DMSO; Sigma-Aldrich, cat. no. 276855) **!** CAUTION: Avoid contact with skin and eyes, and avoid inhalation. Wear protective gloves and clothing. Use it in a chemical fume hood.
- Triethylammonium bicarbonate buffer (TEAB; 1 M, Fluka, cat. no. 17902)
- 4-(2-hydroxyethyl)-1-piperazineethanesulfonic acid buffer (HEPES buffer; Merk, cat. no. 1003378397)
- Tris(2-carboxyethyl)phosphine hydrochloride solution (TCEP; pH 7, 0.5 M, Sigma-Aldrich, cat. no. 646547) **!** CAUTION: Avoid contact with skin and eyes, and avoid inhalation. Wear protective gloves and clothing. Use it in a chemical fume hood.
- Lysyl endoproteinase (Lys-C; Wako Chemicals, cat. no. 129-02541)
- Modified trypsin, in-house synthesized ▴ CRITICAL: We use in-house modified trypsin [60]. Trypsin can be made in advance and stored at -20°C for several months. Alternatively, Promega-sequencing grade modified trypsin can be used.
- Acetonitrile (ACN, HPLC LC-MS grade; VWR Chemicals, cat. no. 83640.290) **!** CAUTION: ACN is toxic and flammable. Avoid contact with skin and eyes, and avoid inhalation. Wear protective gloves and clothing. Use it in a chemical fume hood.
- Tandem Mass Tag Systems (TMTpro 16-plex; Thermo Fisher, cat. no. A44520) ▲ CRITICAL: Although we use a TMTpro 16plex, other multiplexes (TMTpro 18 or 32plex or iTRAQ) can be used if the total protein level in all channels is at least 1 mg.
- Milli-Q Water
- Ammonium bicarbonate (NH_4_HCO_3_; Sigma-Aldrich, cat. no. A6141)
- Trifluoroacetic acid (TFA; Sigma-Aldrich, cat. no. S6046378) **!** CAUTION: TFA is a highly corrosive acid. Avoid contact with skin or eyes, and avoid inhalation. Wear protective gloves and clothing. Use it in a chemical fume hood.
- Glycolic acid (Fluka, cat. no. 50590) **!** CAUTION: Glycolic acid is a corrosive acid. Avoid contact with skin and eyes, and avoid inhalation. Wear protective gloves and clothing. Use it in a chemical fume hood.
- Titansphere TiO_2_ beads (GL Sciences Inc., 5020-75000) **!** CAUTION: Avoid contact with eyes, and wear gloves and eye protection.
- PNGase F (glycerol-free) (New England Biolabs, cat. no. P0705L)
- Sialidase A (Agilent Technologies, cat. no. GK80040)
- Dithiothreitol (DTT; Sigma-Aldrich, cat. no. D9162) **!** CAUTION: DTT is a health hazard compound. Avoid contact with skin and eyes, and avoid inhalation. Wear protective gloves and clothing. Use it in a chemical fume hood.
- Iodoacetamide (IAA; Sigma-Aldrich, cat. no. l1149-25G) **!** CAUTION: IAA is a health hazard compound. Avoid contact with skin and eyes, and avoid inhalation. Wear protective gloves and clothing. Use it in a chemical fume hood.
- Ammonia solution (25%, Sigma-Aldrich, cat. no. 1.05428.0500) **!** CAUTION: Avoid contact with skin and eyes, and avoid inhalation. Wear protective gloves and clothing. Use it in a chemical fume hood.
- Methanol (MeOH; Sigma Aldrich, cat. no. 34860) **!** CAUTION: MeOH is highly toxic and flammable. Avoid contact with skin or eyes, and avoid inhalation. Wear protective gloves and clothing. Use it in a chemical fume hood.
- OLIGO^TM^ R3 reversed-phase resin (Thermo Fisher Scientific, 1-1339-03)
- Sodium hydroxide (NaOH; Sigma Aldrich, cat. no. 415413) **!** CAUTION: Sodium hydroxide is a corrosive basic compound. Avoid contact with skin or eyes, and avoid inhalation. Wear protective gloves and clothing. Use it in a chemical fume hood.
- Agarose, S3 high-capacity acyl-rac capture beads (NANOCS Inc, cat. no. AR-S3-2)
- Hydrochloric Acid (HCl; Merck, cat. no. 1003180500) **!** CAUTION: Hydrogen chloride is a corrosive acid. Protect from direct sunlight and heating sources, as heating can lead to explosion. Avoid contact with skin or eyes, and avoid inhalation. Wear protective gloves and clothing. Use it in a chemical fume hood.
- PTMScan Acetyl-Lysine Motif [Ac-K] Immunoaffinity Beads (Cell Signaling Technology, cat. no. #13362)
- PTMScan IAP Buffer (10X) (Cell Signaling Technology, cat. no. #9993)
- Ammonium formate (Fluka, cat. no. 14266) **!** CAUTION: Avoid contact with skin and eyes, and avoid inhalation. Wear protective gloves and clothing. Use it in a chemical fume hood.

### EQUIPMENT

- Micro tube protein low binding (Sarstedt, cat. no. 72.706.600)
- pH meter
- Sonicator UTR200 (QSonica)
- Water bath
- Centrifuge (capable of centrifuging up to 20,000 xg)
- Nanophotometer N60 (Implen)
- 10 kDa cutoff Amicon Ultra Centrifugal filters (Sigma-Aldrich, cat. no. UFC201024)
- SpeedVac vacuum concentrator (Thermo Scientific)
- Amber light protective Eppendorf tubes
- Vortex mixer
- Rotator (ELMI Intelli-Mixer™ RM-2L, Elmi)
- Oven (capable of heating up to 37°C)
- Shaker (VXR basic Vibrax®, IKA)
- Table centrifuge (microcentrifuge)
- Empore^TM^ SPE disk C8 (Sigma Aldrich, cat. no. 66882-U)
- 30 mg HLB cartridge (Waters, UK)
- Falcon tube (15 mL, Greiner, cat. no. 188261-N)
- Plastic syringe (1 mL, Becton Dickinson, cat. no AB-228000)
- Empore^TM^ SPE disk C18 (Sigma Aldrich, cat. no. 66883-U)
- MobiSpin column filter, 10 µm (MoBiTec Molecular Biology, Germany)
- Low-binding 96-well plates (Corning, cat. no. PCR-96-FS-C)
- Dionex Ultimate 3000 HPLC system (Thermo Fisher Scientific, USA) for High pH Reversed Phase fractionation.
- In-house-made analytical column 22 cm, 100 μm inner diameter, C18 resin column (Reprosil 1.9 µm, Dr. Maisch, Ammerbuch-Entringen, Germany) for non-modified peptides, RmCys peptides, phosphopeptides, and SIA peptides
- In-house-made pre-column (3 cm, 100 μm inner diameter packed with Reprosil-Pur 120 C18-AQ, 5 μm (Dr. Maisch GmbH)) for lysine acetylated peptides and deglycopeptides
- In-house-made pulled emitter analytical column (18 cm, 75 μm ID packed with Reprosil-Pur 120 C18-AQ, 3 μm (Dr. Maisch GmbH)) for lysine acetylated peptides and deglycopeptides
- LC system ▴ CRITICAL: We use an Easy-nLC 1200 (Thermo Scientific), but any nanoLC system can be used.
- Mass spectrometer for LC-MS/MS analysis (Orbitrap Eclipse Tribrid; Orbitrap Exploris 480; Orbitrap Fusion Lumos, Thermo Fisher Scientific) ▴ CRITICAL: Although we used these mass spectrometers, other Orbitrap systems can be used, considering that high resolution in the low mass range and relatively fast scanning is required.
- Column oven/heater, in-house made
- (Optional) Vanquish Horizon UPLC (Thermo Fisher Scientific, Germering, Germany) for metabolomics
- (Optional) ZORBAX pre-column (5 cm, 2.1 mm ID packed with Eclipse Plus C18, 1.8 µm particle size (Agilent Technologies, Santa Clara, CA, USA)) for metabolomics
- (Optional) ZORBAX analytical column (15 cm, 2.1 mm ID packed with Eclipse Plus C18, 1.8 µm particle size (Agilent Technologies, Santa Clara, CA, USA)) for metabolomics SOFTWARE
- Proteome Discoverer (Thermo Scientific, version 2.5.0.400) is used for proteomics/ PTMomics. ▲ CRITICAL: Although we use Proteome Discoverer, any other protein/peptide identification and quantification software can be used if they allow for identification and quantification of TMT 16-plex reagents.
- (Optional) MZmine (version 3.9) is used for metabolomics
- (Optional) MetaboLink (https://computproteomics.bmb.sdu.dk/Metabolomics/) is used for metabolomics

### REAGENT SETUP

#### Samples

The IMR90-4 induced pluripotent stem cell line was purchased from WiCell and used for generating cerebral brain organoids used in this study [55]. The pluripotent stem cells were maintained in feeder-free conditions on Matrigel coated (GFR, Corning) cell culture dishes using mTESR1 medium (StemCell Technologies) at 37°C in 5% CO_2_ with daily medium change. Unguided neural organoids (CBOs) were generated using the commercially available STEMdiff™ Cerebral Organoid Kit (StemCell Technologies) with a few modifications as described in [55]. On days 130, 140, 150, and 160, CBOs were washed in 50 mM ammonium acetate (pH 7.5) and fast frozen in dry ice. Samples were stored at -70°C until further processing.

#### Lysis buffer

Lysis buffer consists of 3% SDC, 10 mM ammonium acetate, pH 7.5, to which PhosSTOP™ must be added just before use. A stock solution of 3% SDC, 10 mM ammonium acetate, pH 7.5 can be made in advance and stored at 4°C for up to one week.

▲ CRITICAL: Ammonium acetate can be replaced by 100 mM HEPES buffer, pH 8.5 if the metabolome is not of interest.

#### Generation of CysPAT (Cysteine-specific Phosphonate Adaptable Tag) reagents

Weigh 5 mg (17.75 μmol) of SIA and 5 mg (32 μmol) of 2-AEP in two separate amber light protective Eppendorf tubes. Then, dissolve SIA in 20 μL of dimethyl sulfoxide (DMSO) and 2-AEP in 400 μL of 100 mM TEAB. Mix the two reagent solutions carefully by adding the 2-AEP solution drop by drop to the SIA solution, while vortexing at low speed. Adjust the pH to 7.5 ∼ 8 by adding 1M TEAB. Then, adjust the final volume to 706 µL using 100 mM TEAB buffer to ensure a final concentration of synthesized CysPAT of 25 mM. Incubate the reagent in the dark for 2 hours at RT with rotation.

▲ CRITICAL: Prepare these solutions fresh just before use because of their unstable nature.
▲ CRITICAL: A CysPAT reagent consists of 1 mg of reagent. Considering a final concentration of 5 mM of CysPAT reagent per sample, for approximately 20 samples, 5x CysPAT reagents (5 mg) are needed.

#### TiO_2_ buffers

- Loading buffer: 1 M glycolic acid in 80% (vol/vol) acetonitrile, 5% (vol/vol) TFA ! CAUTION: TFA is a highly corrosive acid. Handle in a fume hood while wearing gloves.
▲ CRITICAL: The concentration of glycolic acid and TFA is critical to obtain high selectivity in the enrichment.
- Washing buffer 1: 80% (vol/vol) acetonitrile, 1% (vol/vol) TFA
- Elution buffer: ammonia solution 25% NH_4_OH in ultrapure water (pH 11.3)

▲ CRITICAL: Prepare these solutions fresh before use to avoid changes in the buffer composition.

#### PTM-scan kit buffer

Prepare a PTMScan IAP Buffer (1X) solution by diluting the PTMScan IAP Buffer (10X) solution 10X with MilliQ water.

#### Buffers for High pH Reversed-Phase fractionation

- Buffer A: 20 mM ammonium formate, pH 9.3
- Buffer B: 80% acetonitrile, 20% buffer A

#### LC-MS/MS buffers

- Buffer A: 0.1% (vol/vol) FA
- Buffer B: 0.1% (vol/vol) FA, 90% (vol/vol) acetonitrile

### EQUIPMENT SETUP

#### Dionex Reversed-Phase High pH

The solvents are given in REAGENT SETUP. The non-modified peptide fraction and the phosphopeptide/SIA peptide fraction are each separated into 20 concatenated fractions, while the RmCys peptide fraction is fractionated into 12 concatenated fractions.

##### 1. 20 concatenated fraction method

**1. 20 concatenated fraction method:**
| Time (min) | %B | Flow rate (μL/ min) |
| --- | --- | --- |
| 15 | 2 (FT collection) | 5 |
| 20 | 10 | 5 |
| 85 | 40 | 5 |
| 117 | 50 | 5 |
| 122 | 95 | 5 |
| 132 | 95 | 5 |
| 133 | 2 | 5 |
| 148.1 | 2 (FT collection) | 5 |

##### 2. 12 concatenated fraction method

**2. 12 concatenated fraction method:**
| Time (min) | %B | Flow rate (μL/ min) |
| --- | --- | --- |
| 15 | 2 (FT collection) | 5 |
| 74 | 50 | 5 |
| 84 | 70 | 5 |
| 89 | 95 | 5 |
| 99 | 95 | 5 |
| 100 | 2 | 5 |
| 116.1 | 2 (FT collection) | 5 |

#### Liquid chromatography

Solvents A and B are given in REAGENT SETUP. These settings are specific to this nano-Easy LC setup. If another LC system is used, it may be necessary to adjust these parameters.

##### 1. Non-modified peptide and RmCys peptide fractions

Non-modified and RmCys peptide fractions are run on a column of 22 cm, 100 μm inner diameter, ReproSil 1.9 µm C18 material, with the following gradient:

| Time (min) | %B | Flow rate (nl/ min) |
| --- | --- | --- |
| 0 | 2 | 300 |
| 100 | 25 | 300 |
| 120 | 40 | 300 |
| 121 | 95 | 300 |

##### 2. Phosphorylated/CysPAT labelled Cys peptide fraction

The phosphorylated/ CysPAT labelled Cys peptide fraction is separated using the same column specified above for non-modified peptide and RmCys peptide fractions, but with the following gradient parameters:

| Time (min) | %B | Flow rate (nl/ min) |
| --- | --- | --- |
| 0 | 2 | 300 |
| 75 | 25 | 300 |
| 90 | 40 | 300 |
| 91 | 95 | 300 |

##### 3. Deglycopeptide and lysine-acetylated peptide fractions

Deglycopeptide and lysine-acetylated peptide fractions are separated using an in-house-made fused silica capillary two-column setup, consisting of a 3 cm pre-column (100 μm inner diameter packed with Reprosil-Pur 120 C18-AQ, 5 μm), and an 18 cm pulled emitter analytical column (75 μm ID packed with Reprosil-Pur 120 C18-AQ, 3 μm). The following gradient is used:

| Time (min) | %B | Flow rate (nl/ min) |
| --- | --- | --- |
| 0 | 2 | 300 |
| 100 | 25 | 300 |
| 120 | 40 | 300 |
| 121 | 95 | 300 |

##### 4. Metabolite samples (Optional)

Metabolite samples are separated using a Vanquish Horizon UPLC equipped with a 5 mm ZORBAX pre-column (2.1 mm inner diameter packed with Eclipse Plus C18, 1.8 µm) and a 15 cm ZORBAX analytical column (2.1 mm inner diameter packed with Eclipse Plus C18, 1.8 µm) kept at 40°C. The following gradient parameters are set (Solvent A: 0.1% formic acid, Solvent B: 90% formic acid):

| Time (min) | %B | Flow rate (µL/<br>min) |
| --- | --- | --- |
| 0 | 3 | 400 |
| 1.5 | 3 | 400 |
| 4.5 | 40 | 400 |
| 7.5 | 95 | 400 |
| 10.1 | 95 | 400 |
| 10.5 | 3 | 400 |
| 14 | 3 | 400 |

#### Mass spectrometer methods

MS settings used for the LC-MS/MS analysis are tabulated below in the different sections. These parameters are specific for analysis on these mass spectrometers. If another MS system is used, it may be necessary to adjust these parameters.

##### 1. Non-modified peptide fractions

For non-modified peptide fractions, an Orbitrap Eclipse Tribrid mass spectrometer is used with the following settings:

▲ CRITICAL: The method described here applies exclusively to the Orbitrap Eclipse Tribrid. In addition, it is important to consider that the use of SPS MS^3^ [61] restricts the dataset and enzyme options in the subsequent database search analysis. Alternatively, the instrument and method described in “*2. Phosphorylated/SIA peptide and RmCys peptide fractions***”** can be used.

| Parameter | Value |
| --- | --- |
| ***Full MS*** |  |
| **Detector** | orbitrap |
| **Polarity** | positive |
| **Scan range m/z** | 350-1600 |
| **Resolution** | 120,000 FWHM |
| **AGC value** | 250% |
| **Max injection time** | 50 ms |
| ***Data-dependent dd-MS<sup>2</sup>*** |  |
| **Detector** | Ion trap |
| **MS/MS cycle time** | 3 sec |
| **Isolation window** | 0.7 Da |
| **Activation type** | CID |
| **Collision energy type** | normalized |
| **CID normalized collision energy (NCE)** | 35% |
| **Turboscan** | on |
| ***SPS-MS<sup>3</sup>*** |  |
| **Detector** | orbitrap |
| **Isolation window** | 2 |
| **N of SPS precursors** | 10 |
| **Activation type** | HCD |
| **TurboTMT** | TMT and TMTpro Reagents |
| **HCD NCE** | 55% |

##### 2. Phosphorylated/SIA peptide and RmCys peptide fractions

For the LC-MS/MS analysis of phosphorylated/SIA peptide and RmCys peptide fractions, an Orbitrap Exploris 480 mass spectrometer is used with the following parameters:

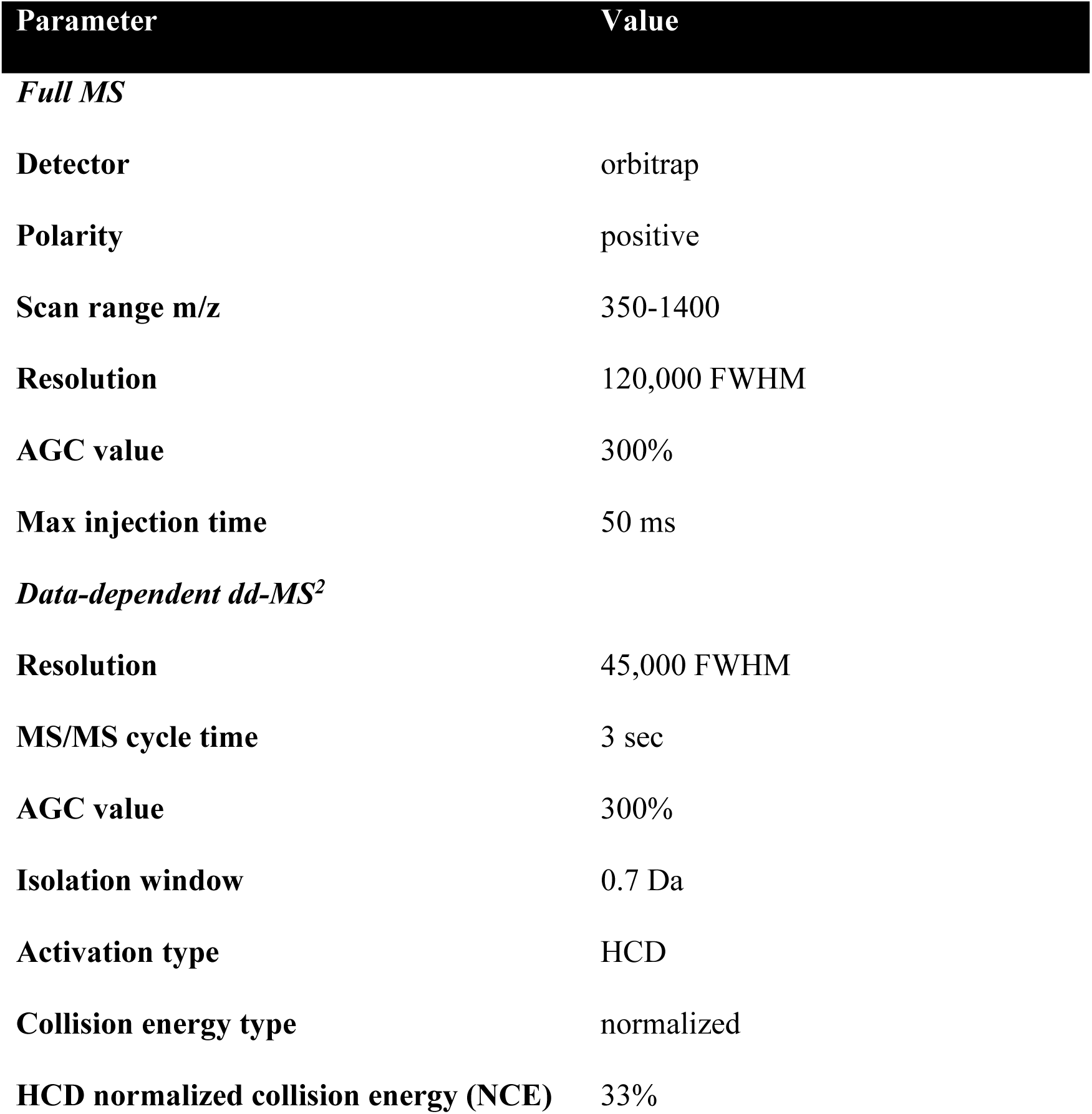
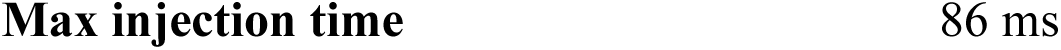

##### 3. Deglycopeptide and lysine-acetylated peptide fractions

An Orbitrap Fusion Lumos mass spectrometer is used to analyze the deglycopeptide and lysine-acetylated peptide fractions with the following parameters:

| Parameter | Value |
| --- | --- |
| *Full MS* |  |
| Detector | orbitrap |
| Polarity | positive |
| Scan range m/z | 350-1200 |
| Resolution | 120,000 FWHM |
| AGC value | 250% |
| Max injection time | 50 ms |
| *Data-dependent dd-MS<sup>2</sup>* |  |
| Detector | orbitrap |
| Resolution | 50,000 FWHM |
| MS/MS cycle time | 3 sec |
| AGC value | 300% |
| Isolation window | 0.7 Da |
| Activation type | HCD |
| Collision energy type | normalized |
| HCD normalized collision energy (NCE) | 34% |
| Max injection time | 86 ms |

##### 4. Metabolites (Optional)

A Q Exactive HF mass spectrometer is used for the LC-MS/MS analysis of the metabolome samples with the following parameters:

▲ CRITICAL: We recommend using a fraction of each individual metabolomic sample for a pooled sample, which is used as a quality control (QC) sample for data filtering and normalization during data processing steps. This sample should be analyzed regularly during sample analysis on the mass spectrometer to correct for a potential change in instrument performance over time.

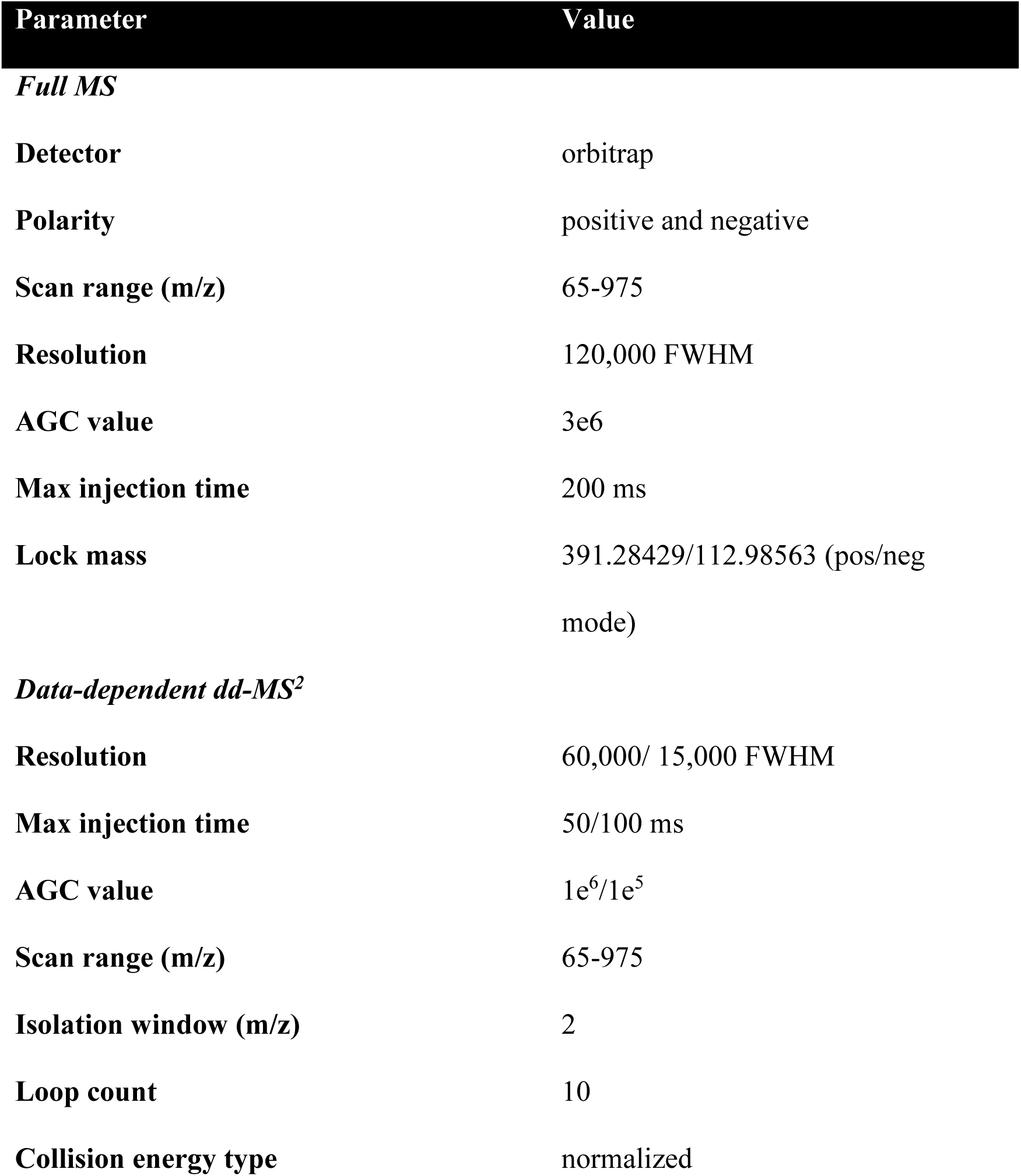
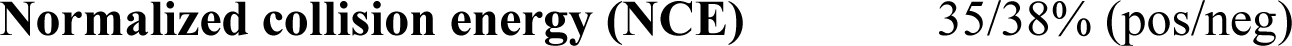

#### Database searching

##### 1. Proteomics and PTMomics

The proteomics and PTMomics data are processed with Proteome Discoverer (Version 2.5.0.400, Thermo Fisher Scientific) and subjected to database searching using the Sequest HT search engine with the following criteria: human proteome as database (version from 2023.06.28), trypsin as enzyme with 2 missed cleavages allowed (up to 4 missed cleavages is allowed for the lysine-acetylated peptides), and TMTpro on the peptide N-terminus and lysine residues as fixed modifications (TMTpro on lysine residues as a dynamic modification for the lysine-acetylated peptides). In addition, for PTMome searches, dynamic modifications included are as follows (summarized in table below): acetylation of lysine residues for the lysine-acetylated peptides, carbamidomethylation of cysteine residues for the RmCys-containing peptides and the depalmitoylated peptides (optional), SIA on cysteine residues and phosphorylation of serine, threonine, and tyrosine residues for the free cysteine- and phosphorylation-containing peptides, and deamidation of asparagine residues for the formerly sialylated N-glycopeptides. 10 ppm and 0.05 Da (0.8 Da if SPS MS^3^ was used (non-modified peptides)) are set respectively as precursor mass tolerance and fragment mass tolerance. Identifications are filtered against a 1% false discovery rate cut-off using the integrated Percolator algorithm [62]. Quantification is performed based on TMT reporter ion intensities measured in MS^2^ scans for PTMomic data and in MS^3^ scans for proteomic data (non-modified peptides).

Specifically, for each dataset, only peptides carrying the following modifications are considered:

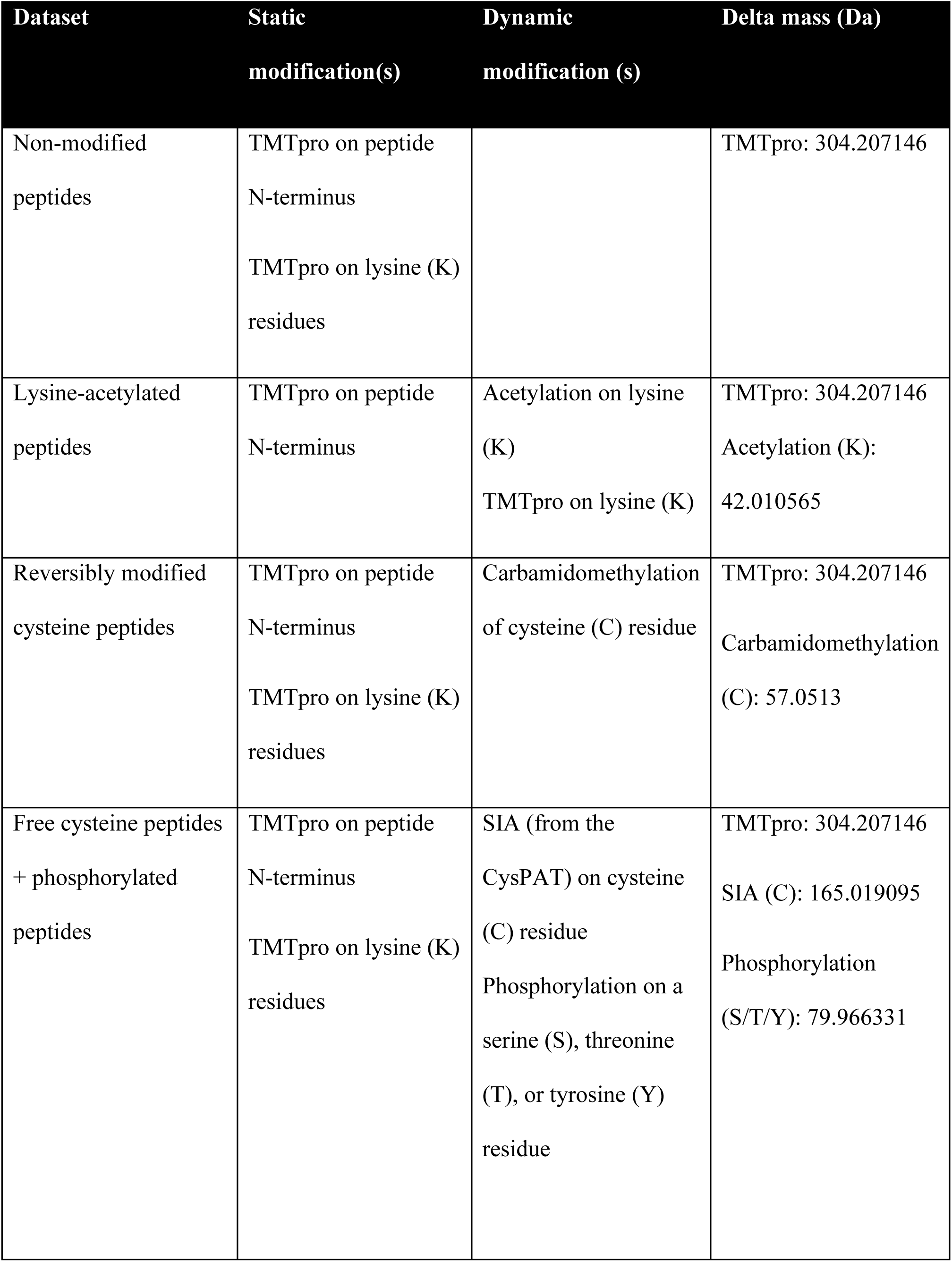
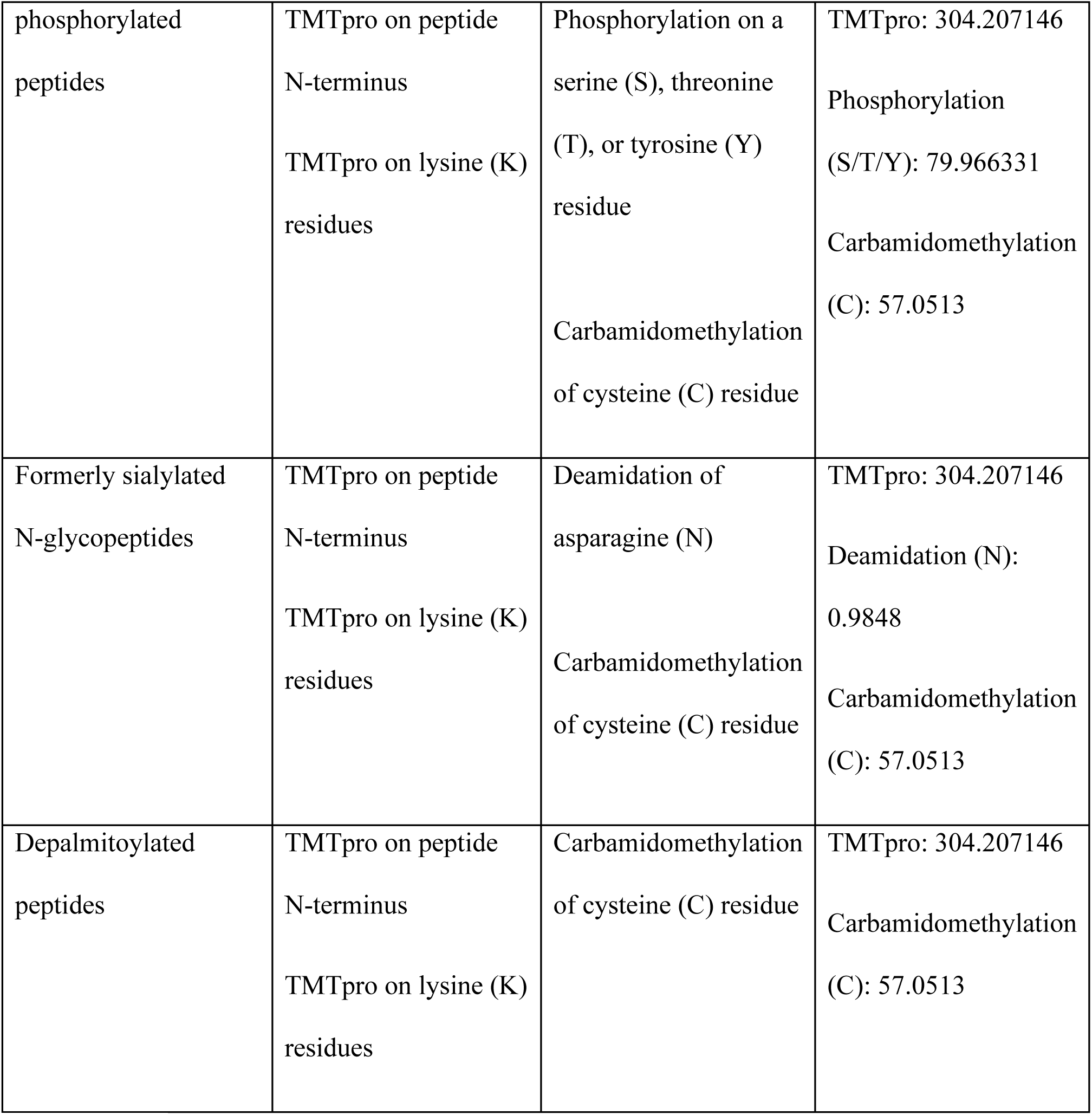

▴ CRITICAL: Deamidation can spontaneously occur during the sample preparation [63]. A further filtering step is recommended: accept only formerly sialylated N-glycopeptides presenting the N-linked glycosylation motif (N-X-S/T/C, X≠P), and known to be N-glycosylated according to the UniProt Knowledgebase.
▲ CRITICAL: To identify CysPAT-labelled free cysteine-containing peptides, the CysPAT modification must be added as a modification to the database used for searching. Add the CysPAT modification using the following settings: Modification = CysPAT, Delta Mass [Da] = 165.019095, Delta Average Mass [Da] = 165.0844, Substitution = C(4) H(8) N O(4) P, Position = Any, Target = C.
▲ CRITICAL: To avoid unnecessary expansion of search space and potential increase of false discovery rates (FDR), variable modifications should be limited to those that are expected to be present in each of the PTM fractions.
▲ CRITICAL: The use of TMT as variable modification in database searching is strongly discouraged, except when searching data from the lysine-acetylation fraction. Testing the TMT labeling efficiency prior to PTM enrichment (step 39) allows the application of TMT as fixed modification, thereby avoiding unnecessary expansion of the search space.
▲ CRITICAL: Several peptide modifications can arise during sample handling, including methionine oxidation and asparagine/glutamine deamidation (N/Q). Spontaneous deamidation of N and Q residues occurs under neutral to slightly basic pH conditions[63]. For this reason, we generally do not recommend including methionine oxidation or deamidation as variable modifications in PTM characterization -unless there is a strong experimental rationale for this, such as including deamidation (N) when searching N-deglycosylated peptides.

##### 2. Metabolomics (Optional)

Metabolomics data are processed with MZmine (version 3.9) [64] for the raw files obtained by positive and negative ion mode separately. The following modules and settings are used to extract ion chromatograms and perform steps of chromatogram deconvolution, deisotoping and alignment across the samples: Mass detection, Chromatogram builder, Local minimum resolver, 13C Isotope feature filter, Join aligner, and Gap filling (Same m/z and RT range gap filler, m/z tolerance: 10 ppm). Feature peaks are annotated in MZmine according to the Metabolomics Standards Initiative (MSI) level 3 using local MS/MS spectra databases of National Institute of Standards and Technology 17 (NIST17), Mass Bank of North America (MoNA). After annotation of feature peaks, the datasets from running in positive and negative ion mode are filtered and normalized according to the common QC samples in MetaboLink (https://computproteomics.bmb.sdu.dk/Metabolomics/) [65]. In MetaboLink, normalization is performed using the Probabilistic Quotient Normalization method, and the positive and negative ion mode datasets are merged using the ‘Merge datasets’ function.

#### Procedure

##### Tissue lysis **●** Timing 6 h for 16 samples

1. Add 300 µL of lysis buffer to the sample.

▲ CRITICAL: Add PhosStop to the lysis buffer just before use.
▲ CRITICAL: The amount of lysis buffer needs to be adjusted based on the size of the tissue/sample.
▲ CRITICAL: For more complex samples (e.g., tissues) it may be necessary to add an additional step of homogenization, such as bead beating, and higher concentration of SDC (5%) in the lysis buffer. Additional time then needs to be allocated for this step.
**?** TROUBLESHOOTING
2. Sonicate at 40% amplitude for 4x 10 sec

▲ CRITICAL: Make sure there is no visible pellet in the sample after the sonication.
▲ CRITICAL: Avoid the formation of foam during the sonication.
**?** TROUBLESHOOTING
3. Heat the sample up for 5 min at 100°C in a water bath.

▲ CRITICAL: Given the high temperature and increasing pressure inside the tube, it is recommended to make a small hole in the lid or open the lid.
**?** TROUBLESHOOTING
4. Centrifuge the samples for 20 min at 20,000 xg at RT to pellet insoluble material.
5. Transfer the supernatant to a new low binding Eppendorf tube.
6. Use around 1 µL of sample to measure the protein concentration using the N60 nanophotometer.
7. Transfer 150 µg protein for each sample to a 10 kDa spin filter.
**?** TROUBLESHOOTING

##### (Optional) Metabolite extraction ● Timing 2 h for 16 samples + vacuum centrifugation

8. After transferring 150 µg protein for each sample to a 10 kDa spin filter, adjust the final volume to 100 µL by adding lysis buffer.
9. Dilute the sample to 200 µL using 10 mM ammonium acetate, pH 7.5. Ensure that the final concentration of SDC is 1%.
10. Centrifuge at 14,000 xg at RT for 30 min (or until the solution has passed through the filter).
11. Transfer the flow-through from the 10 kDa spin filter to a new low-binding Eppendorf tube.

▴ CRITICAL: Proteins are retained on top of the 10 kDa spin filter. Save the filter and proceed with step 17 after preparing the CysPAT reagent.
▴ CRITICAL: If the experiment consists of a larger number of samples, it is recommended to transfer the 10 kDa flow-through to a non-dripping 0.45 µm filter plate instead of Eppendorf tubes.
12. To the 10 kDa flow-through, add 6 µL 100% formic acid (final concentration of 2%) to precipitate SDC and mix it very well.

▴ CRITICAL: Lipids will precipitate with SDC and therefore will not be analyzed in the metabolomics sample.
13. Centrifuge the samples at 20,000 xg at RT for 20 min (or until the solution has passed through the filter) to remove SDC from the small molecule metabolites.
14. Transfer 200-250 µL from the flow-through to a new low-binding 96-well plate.

▲ CRITICAL: It is important to transfer the same volume from all samples to enable normalization of metabolite abundance to the protein content of the sample.
15. Dry down the metabolite sample by vacuum centrifugation and store it at -20°C for metabolomic analysis.

##### CysPAT reagent preparation ● Timing 30 minutes + 2 hours incubation time

▲ CRITICAL: Prepare these solutions just before use because of their unstable nature. Use the 2 hours of incubation to perform tissue lysis or metabolite extraction.
▲ CRITICAL: 1x CysPAT reagent consists of 1 mg of reagent. For 20 samples 5x CysPAT reagents (5 mg) are needed.
16. Prepare the CysPAT reagents according to *Generation of CysPAT (Cysteine-specific Phosphonate Adaptable Tag) reagents* as described in the REAGENT SETUP section.

##### CysPAT labeling of free cysteines ● Timing 3.5 hours

17. Add 100 µL 100 mM HEPES buffer containing 1% SDC, pH 8.5 to the proteins on top of the 10 kDa spin filter.
18. When CysPAT is ready, add 25 µL of CysPAT reagents (final concentration around 5 mM) to each sample and mix it carefully.

▲ CRITICAL: The volumes of samples and CysPAT can be changed, however it is advisable to use a final concentration of CysPAT reagent equal to approximately 5 mM.
19. Incubate at 37°C for 1 hour.
20. Centrifuge the sample at 11,000 xg for 30 min at RT to reduce the volume and remove excess CysPAT reagent.

▴ CRITICAL: If you are not performing metabolomics, transfer the solution to a 10 kDa spin filter tube before centrifugation.
▲ CRITICAL: It is possible to dilute the sample with 100 µL 50 mM HEPES buffer before centrifugation to help the solution pass through the filter.
21. Wash the proteins on top of the filter using 400 µL of 100 mM HEPES buffer containing 0.2% SDC, pH 8.5 and centrifuge at 11,000 xg for 30 min at RT. Perform this wash twice.
**?** TROUBLESHOOTING

##### Protein reduction and digestion ● Timing 3.5 hours + overnight digestion

22. Add 100 µL of 50 mM HEPES buffer containing 1% SDC and 3 mM TCEP. Incubate at RT for 30 min. **?** TROUBLESHOOTING
23. Digest the proteins by adding Endoproteinase Lys-C (0.04 units (AU) per 1 mg of protein), and incubate for 1 hour at 37°C.
24. Add trypsin to the solution in an enzyme/substrate ratio of between 1:50 to 1:20 (w/w) and incubate at 37°C over-night.

▲ CRITICAL: It is possible to reduce the incubation time for trypsin to 4 hours.
25. After the incubation, add trypsin in an enzyme/substrate ratio of 1:100 (w/w), and continue the incubation for 1 hour at 37°C.
26. Transfer the peptide solution from the top of the filter to a new low-binding tube.
27. Wash the filter using 30 µL 30% acetonitrile and transfer the wash solution to the low-binding tube with the peptide solution.

▪ PAUSE POINT: Samples can be frozen and stored at −20°C.

##### (Optional) Evaluation of the digestion quality ● Timing 30 min + LC-MS/MS

28. Take out 2 µL of the sample.

▲ CRITICAL: There is no need to check the digestion quality of all samples; we recommend instead performing it only on 3-4 samples.
29. Add 10 µL of 3% formic acid and vortex it to precipitate the SDC.
30. Centrifuge at 14,000 xg for 20 min at RT to pellet the SDC.
31. Transfer 5 µL of the supernatant to a 96-well plate.
32. Analyze the sample on the LC-MS/MS instrument to evaluate the digestion efficiency using the parameters described in the EQUIPMENT SETUP section.

▲ CRITICAL: (Optional) It is possible to check the peptide quantification of the samples, running 3-4 HeLa standards with the same setup and the same settings to make a standard curve. We usually run 3 HeLa standards of 100, 200, and 300 ng.
**?** TROUBLESHOOTING

##### TMT labeling ● Timing 3 hours + vacuum centrifugation + LC-MS/MS (Optional)

33. Dry down the digested peptide solution by vacuum centrifugation.
34. Resolubilize the peptides in 100 µL 50 mM HEPES buffer, pH 8.5.

▲ CRITICAL: It is essential that SDC is resolubilized. If SDC is difficult to solubilize, leave the sample on a shaker for 30 min before proceeding.
35. Reconstitute each 0.5 mg TMT reagent in 20 µL of acetonitrile, allow it to sit for 10 min with occasional vortexing. Briefly spin down the TMT reagent using the table centrifuge to gather the solution at the bottom.
36. Add each TMT reagent to the corresponding sample and mix it well.
37. Adjust the pH to 8-8.5 using 1M TEAB if needed.

▲ CRITICAL: A pH outside this range negatively affects the labeling efficiency of TMT.
38. Incubate for 1.5 hours at RT.
39. After the incubation, take an aliquot of 1 µL of each sample and add them together into 30 µL H_2_O, to create a QC test and check the labeling. After mixing add 2% (v/v) formic acid and mix well. Centrifuge at 14,000 xg for 10 min to pellet the SDC and analyse the QC test sample by LC-MS/MS.

▲ CRITICAL: If the TMT incorporation is lower than 90%, perform an additional incubation with TMT reagents, and check again the labeling. **?** TROUBLESHOOTING
40. Store the rest of the samples at 4°C while testing the labelling efficiency.

▲ CRITICAL: Do not quench and mix the samples before assessing the TMT labeling quality.
▪ PAUSE POINT: Samples can be stored at −20°C for up to 2 weeks.
41. Mix the TMT samples according to the QC test analysis. Then, add 10 µL of 1 M ammonium bicarbonate and incubate for 15 min at RT to quench the TMT reaction.

▲ CRITICAL: Do not quench the samples using hydroxylamine, if you aim at performing S-palmitoylation enrichment.

##### Precipitation of SDC ● Timing 30 min + vacuum centrifugation

42. Acidify the TMT mix sample with 2% (v/v) of formic acid and vortex.
43. Centrifuge at 20,000 xg for 15 min to pellet SDC.
44. Transfer the supernatant to a new low binding tube.

▲ CRITICAL: (Optional) It is possible to resolubilize the SDC pellet with 500 mL of 50 mM HEPES buffer and adjust the pH to 8-9. Then, precipitate again with 2% (v/v) formic acid, centrifuge as above and pool the new supernatant with the first supernatant. This optional step gives more peptides to continue with.
▲ CRITICAL: (Optional) It is possible to save the SDC pellet at –20°C and use it to enrich S-palmitoylated peptides as described elsewhere [52].
45. Using a vacuum centrifuge, lower the volume of the solution to maximum 150 µL.

##### TiO_2_ enrichment ● Timing 70 min + vacuum centrifugation

46. Make the sample up to the TiO_2_ loading buffer composition (80% acetonitrile, 5% TFA and 1 M glycolic acid).

▲ CRITICAL: If the sample has a volume of 100 µL, add 50 µL of milli-Q water, 50 µL of 100% TFA, 800 µL of acetonitrile and 76 mg of glycolic acid.
! CAUTION: TFA is a highly corrosive acid. Avoid contact with skin or eyes and avoid inhalation. Handle in a chemical fume hood and wear protective gloves.
▲ CRITICAL: The concentration of glycolic acid and TFA is critical to obtain high selectivity in the enrichment.
47. Add the TiO_2_ beads in a beads-to-peptides ratio of 1 mg per 100 µg of peptides.
48. Place the sample on the shaker at the highest shaking speed (approx. 2,000 rpm) for 15 min at RT.
49. Centrifuge using a table centrifuge for a maximum of 15 sec to pellet the TiO_2_ beads.
50. Transfer the supernatant to another low-binding Eppendorf tube.
51. Use the supernatant for a second incubation using half of the amount of TiO_2_ beads used in the first incubation (beads-to-peptides ratio of 0.5 mg per 100 µg).
52. Place the sample on the shaker at the highest shaking speed (approx. 2,000 rpm) for 10 min at RT.
53. Centrifuge using a table centrifuge for a maximum of 15 sec to pellet the TiO_2_ beads.
54. Transfer the supernatant, containing non-modified peptides (i.e. faction without phosphopeptides, sialylated N-glycopeptides, and/or CysPAT-labelled free cysteine containing peptides), to a new low-binding Eppendorf tube, hereafter referred to as “*TiO_2_ FT sample”*.
**?** TROUBLESHOOTING
55. Use 200 µL of washing buffer 1 to combine the TiO_2_ beads from the two incubations and transfer them to a new low-binding Eppendorf tube.

▲ CRITICAL: The transfer to a new low-binding Eppendorf tube is performed to prevent peptides, which may be attached to the tube surfaces, from co-eluting in the last elution step (steps 74-79).
56. Vortex the solution for 10 sec and then, using a table centrifuge, pellet the beads.
57. Collect the supernatant in the *TiO_2_ FT sample*.

**?** TROUBLESHOOTING
58. Wash the beads with 100 µL of washing buffer 1. Mix it for 10 sec and then, using a table centrifuge, centrifuge to pellet the beads.
59. Collect the supernatant in the *TiO_2_ FT sample*.
60. Dry down the *TiO_2_ FT sample* by vacuum centrifugation.

##### Deglycosylation and enrichment of deglycopeptides ● Timing 2 hours + overnight deglycosylation + vacuum centrifugation

61. Dry the TiO_2_ beads for 5-10 min using the vacuum centrifuge.

▲ CRITICAL Make sure that the beads are completely dry. The presence of washing buffer 1 will compromise the next steps.
62. Redissolve the TiO_2_ beads in 150 µL of 100 mM HEPES buffer (pH 8).
63. Adjust the pH to 8.
64. Add 1 µL of PNGaseF and 0.5 µL of Sialidase A.
65. Incubate overnight at 37 °C.
66. After incubation, add 10 mM DTT and incubate at RT for 30 min.
67. After incubation, add 20 mM IAA and incubate for 30 min in the dark at RT.
68. This step is to alkylate all reduced cysteine residues situated on the enriched peptides.
69. After incubation, add 10 µL of 100% TFA and 500 µL of acetonitrile and mix.
70. Incubate with rotation for 10 min at RT to allow the phosphopeptides and SIA peptides to reattach to the TiO_2_ beads.
71. Pellet the TiO_2_ beads by centrifuging the sample at 14,000 xg.
72. Collect the deglycosylated peptides by transferring the supernatant to a new low-binding Eppendorf tube, named *Deglycopeptides*.
73. Lyophilize the deglycopeptide sample using a vacuum centrifuge and store it for Oligo R3 RP desalting.

▪ PAUSE POINT: Samples can be frozen and stored at −20°C.

##### Collection of phosphopeptides and CysPAT-peptides ● Timing 1 hour + vacuum centrifugation

74. Dry the TiO_2_ beads for 5 min by vacuum centrifugation.
75. Add 150 µL of elution buffer and mix well.

▴ CRITICAL: The elution buffer needs to be freshly prepared before use.
▴ CRITICAL: For a more efficient elution, it is possible to check and, if necessary, adjust the pH to around 11.
76. Incubate the solution on a shaker for 15 min to allow an efficient elution of phosphopeptides and CysPAT*-*peptides.
77. Centrifuge the solution for 1 min (highest speed) to pellet the TiO_2_ beads.
78. Pass the supernatant through a p200 pipette tip with a C8 disc (C8 stage tip) to capture the TiO_2_ beads over the filter. Recover the liquid in a low-binding Eppendorf tube labelled *Phosphopeptides/CysPAT-peptides*.
**?** TROUBLESHOOTING
79. Wash the TiO_2_ beads with 50 µL of 5% ammonia water/50% acetonitrile (pH 11), mix it and centrifuge.
80. Pass the supernatant through the same C8 filter as above and collect the solution into the previous elution (*Phosphopeptides/CysPAT-peptides)*.

▲ CRITICAL: This second TiO_2_ bead elution will also co-elute peptides that could be retained on the C8 filter.
**?** TROUBLESHOOTING
81. Dry the *Phosphopeptides/CysPAT-peptides* in the vacuum centrifuge prior to high pH reversed-phase fractionation.

▲ CRITICAL: Be aware that CysPAT labeling of free cysteine residues and subsequent co-enrichment and analysis of CysPAT- and phosphopeptides can decrease the amount of identified phosphopeptides due to their lower stoichiometry. To increase the number of identified phosphopeptides, more fractions during high pH RP fractionation can be collected.
▲ CRITICAL: (Optional) It is possible to enrich for tyrosine phosphorylated peptides using phosphotyrosine antibodies or SH2 superbinder beads after this step. However, be aware that the efficiency of phosphotyrosine enrichment depends on the amount of starting material. We advise using at least 1.5 mg of protein/peptide if introducing this step.

##### Desalting of *TiO_2_ FT fraction* ● Timing 1.5 hours + vacuum centrifugation

82. Resuspend the *TiO_2_ FT fraction* in 1 mL of 0.1% TFA.
83. Adjust the pH to 2 or lower.
84. Activate an HLB column by passing 2 mL of MeOH followed by 2 mL of 100% ACN.

▲ CRITICAL: Choose the appropriate HLB column based on the amount of material expected in the sample.
E.g. the capacity of HLB column is approximately 10% of its mass. So, having as starting material 16 samples, each containing 150 µg of proteins, we expect not more than 2.4 mg of peptides. Thus, a 30 mg HLB column is appropriate.
**!** CAUTION: MeOH is highly toxic. Work in a chemical fume hood, wearing gloves and protective clothing.
**!** CAUTION: ACN is toxic. Handle it with protective gloves and in a chemical fume hood.
85. Equilibrate the column with 3 mL of 0.1% TFA

▲ CRITICAL: Make sure to not dry the column from now on.
86. Slowly load the sample onto the column.
87. Collect the flow-through in a 15 mL Falcon tube.
88. Reload the sample and pass it through the column a second time, collecting the flow-through.
89. Wash the column with 3 mL of 0.1% TFA and collect the wash.
90. Wash the column with 1 mL of H_2_O and collect the wash.
91. Slowly elute the peptides using 750 µL of 50% ACN, in H_2_O and collect the elute in a low-binding tube.
92. Repeat step 90 using 750 µL of 70% ACN, in H_2_O. Collect the eluate in the same low-binding tube of the previous step.
93. Dry the sample in the vacuum centrifuge.

##### Oligo R3 reversed-phase desalting ● Timing 1 hour

94. Cut the wide end of a p200 pipette tip to fit a plastic syringe.

▲ CRITICAL: Make sure that the syringe makes the vacuum when placed on the tip.
95. Place a small disk of a 3M Empore C18 at the end of the tip.
96. Add Oligo R3 RP material and pack a 1-1.5 cm column by gently flowing air using the syringe.

▲ CRITICAL: The length of the R3 column must be determined based on the amount of peptide expected in the solution. The peptide binding capacity of R3 resin is 1-5% of its weight. A 1 cm column weighs approximately 4.8 mg.
97. Activate the column by flowing 60 µL of ACN through.

▲ CRITICAL: Make sure to not dry the column from now on.
98. Equilibrate the column using 60 µL of 0.1% TFA.
99. Load slowly the acidified peptide sample.

▲ CRITICAL: Make sure the pH of the sample solution is < 2.
100. Wash the column using 60 µL of 0.1% TFA

▲ CRITICAL: Flowthrough collection is highly advisable.
101. Elute the peptides using 60 µL of 65% ACN in H_2_O and collect them in a low binding Eppendorf tube.

##### Purification of reversibly modified cysteine peptides ● Timing 4.5 hours

102. Redissolve the lyophilized peptides from the desalted *TiO_2_ FT* fraction in 50 µL 500 mM HEPES buffer.
103. Adjust the pH to 8 using NaOH if needed.
104. Add TCEP to a final concentration of 5 mM.

▲ CRITICAL: It is essential to keep the concentration of TCEP at maximum 5 mM to not prevent the subsequent binding between peptides and the S3 beads.
105. Incubate at 37°C for 1 hour to reduce again the peptides and prevent disulfide bindings between peptides.
106. During the incubation, take 300 µL of Agarose, S3 high-capacity acyl-rac capture beads.
107. Wash the beads twice using 100 mM HEPES (pH 8).
108. After the incubation with TCEP, dilute rapidly the peptide solution in H_2_O to a final volume of 1.2 mL. Then, add the S3 beads immediately after.
109. Mix well and incubate for 1 h at RT on rotation.
110.After the incubation, pellet the beads by gentle centrifugation using a table centrifuge.
111. Transfer the supernatant to a new low-binding Eppendorf tube, labelled “*S3 FT fraction*”.
112. Add 200 µL of 100 mM HEPES (pH 8), mix shortly, and centrifuge to pellet the beads.
113. Transfer the supernatant to the “*S3 FT fraction*” tube.
114. Solubilize the beads in 500 µL of 100 mM HEPES (pH 8) and mix gently.
115. Transfer the beads to a MobiSpin column filter.
116. Place the MobiSpin column filter in an Eppendorf tube with the lid cut off.
117. Wash the beads twice with 300 µL of water.
118. Block the MobiSpin column filter at the end.
119. Add to the “dry” beads on the top of the spin filter 300 µL of 100 mM HEPES (pH 8) containing 20 mM DTT to release the cysteine-containing peptides from the beads.
120. Mix to get the beads into solution using a pipette tip where the end has been cut off to create a wider bore that enables mixing.
121. Put the lid on the MobiSpin column filter and incubate at 37 for 30 min.
122. Transfer the MobiSpin column filter to a new Eppendorf tube. Then, release partially the lid of the MobiSpin column filter, and remove the “block” previously placed at the end of the column filter.
123. Centrifuge and collect the solution containing the reversibly modified cysteine-containing peptides in a low-binding Eppendorf tube, labelled “*RmCys peptides”*.
124. Wash the beads using 100 µL of 100 mM HEPES, 20 mM DTT (pH 8), and collect the solution in the “*RmCys peptides”* Eppendorf tube.
125. Add 45 mM iodoacetamide to the solution.

▴ CRITICAL: The amount of IAA must be at least double the amount of DTT already present in the solution.
126. Incubate for 30 min in the dark at RT.
127. After incubation, acidify with FA
128. Perform an R3 purification as described in *Oligo R3 reversed-phase desalting* (steps 93-100) and then proceed to the high pH reversed-phase fractionation.

##### Enrichment of lysine acetylated peptides ● Timing 3 hours + vacuum centrifugation

129. Adjust the pH of “*S3 FT fraction*” to pH 7.2 using 1M HCl.
130. Wash 30 µL of the Lys-Ac-beads with washing buffer (IAP buffer) from the PTMScan kit.
131. Add the Lys-Ac-beads to the “*S3 FT fraction*”
132. Incubate for 2 hours at RT under gentle rotation.
133. Gently centrifuge the sample to pellet the beads and transfer the supernatant to a low-binding Eppendorf tube, labeled *“non-modified peptides”*.

▴ CRITICAL: (Optional) It is possible to also perform other antibody-based enrichment of PTM-containing peptides, e.g. O-GlcNAcylated peptides, using the “*non-modified peptides*” sample.
134. Wash the beads 3 times using 200 µL of washing buffer (IAP buffer) from the PTMScan kit.
135. Wash the beads twice using 200 µL of H_2_O.
136. Elute the lysine acetylated peptides by incubating the beads with 150 µL of 0.15% TFA for 20 min at RT under mild shaking.
137. Centrifuge the beads and transfer the supernatant to a new low-binding Eppendorf tube labeled *“LysAc peptides”*.
138. Wash the beads, incubating them with 100 µL of 0.15% TFA for 5 min.
139. Centrifuge the beads and combine the supernatant with previous supernatant.
140. Use an R3 column to desalt the sample as previously described in *Oligo R3 reversed-phase desalting* (steps 93-100)
141. Dry the eluate by vacuum centrifugation.

##### High pH RP fractionation of the non-modified peptides, phosphopeptides/CysPAT peptides and RmCys peptides ● Timing 2.5 hours for 20 fraction method; 2 hours for 12 fraction method + vacuum centrifugation

142. Resuspend the lyophilized fractions in 32 µL of High pH buffer B (pH 9).
143. Vortex well the samples and centrifuge at 14,000 xg for 10 min at RT.
144. Transfer 30 µL of sample into a microtiter plate.
145. Fractionate the samples using the methods described in EQUIPMENT SETUP section, under *Dionex reversed phase High pH*.
**?** TROUBLESHOOTING
146. Dry the fractions contained in the microtiter plate by vacuum centrifugation.

##### LC-MS/MS analysis ● Timing >1 day

147. Reconstituted the lyophilized fractions in MS/MS solvent A.
148. Analyze the samples by LC-MS/MS following the methods described in the EQUIPMENT SETUP section.
**?** TROUBLESHOOTING

##### Data analysis ● Timing >1 day

149. Search the LC-MS/MS raw data files (*non-modified peptides, lysine acetylated peptides, RmCys peptides, phosphopeptides/CysPAT-peptides,* and *deglycopeptides*) in Proteome Discover, using an appropriate database and the search parameters as described in the EQUIPMENT SETUP section.

▲ CRITICAL: Any other MS protein/peptide identification and quantification software can be used with similar settings.
150. (Optional): Process the LC-MS/MS raw data files (*metabolomics*) in MZmine and subsequent normalization and data set merging in MetaboLink as described in the EQUIPMENT SETUP section.

▲ CRITICAL: Other similar software(s) can be used with similar settings.

#### Timing

Steps 1-7, 6 h

Steps 8-15 (Optional), 2 h + vacuum centrifugation

Steps 16, 30 min + 2 h incubation

Steps 17-21, 3.5 h

Steps 22-27, 3.5 h + overnight digestion

Steps 28-32 (Optional), 30 min + LC-MS/MS

Steps 33-41, 3 h + vacuum centrifugation + LC-MS/MS (Optional)

Steps 42-45, 30 min + vacuum centrifugation

Steps 46-60, 70 min + vacuum centrifugation

Steps 71-72, 2 h + overnight deglycosylation + vacuum centrifugation

Steps 73-80, 1h + vacuum centrifugation

Steps 81-92, 1.5 h + vacuum centrifugation

Steps 93-100, 1 h

Steps 101-127, 4.5 h

Steps 128-140, 3 h + vacuum centrifugation

Steps 141-145, 2.5 h (20 fractions) and/or 2 h (12 fractions) + vacuum centrifugation

Steps 146-147, > 1 d

Steps 148-149> 1 d

#### Troubleshooting

Troubleshooting is summarized in the following table.

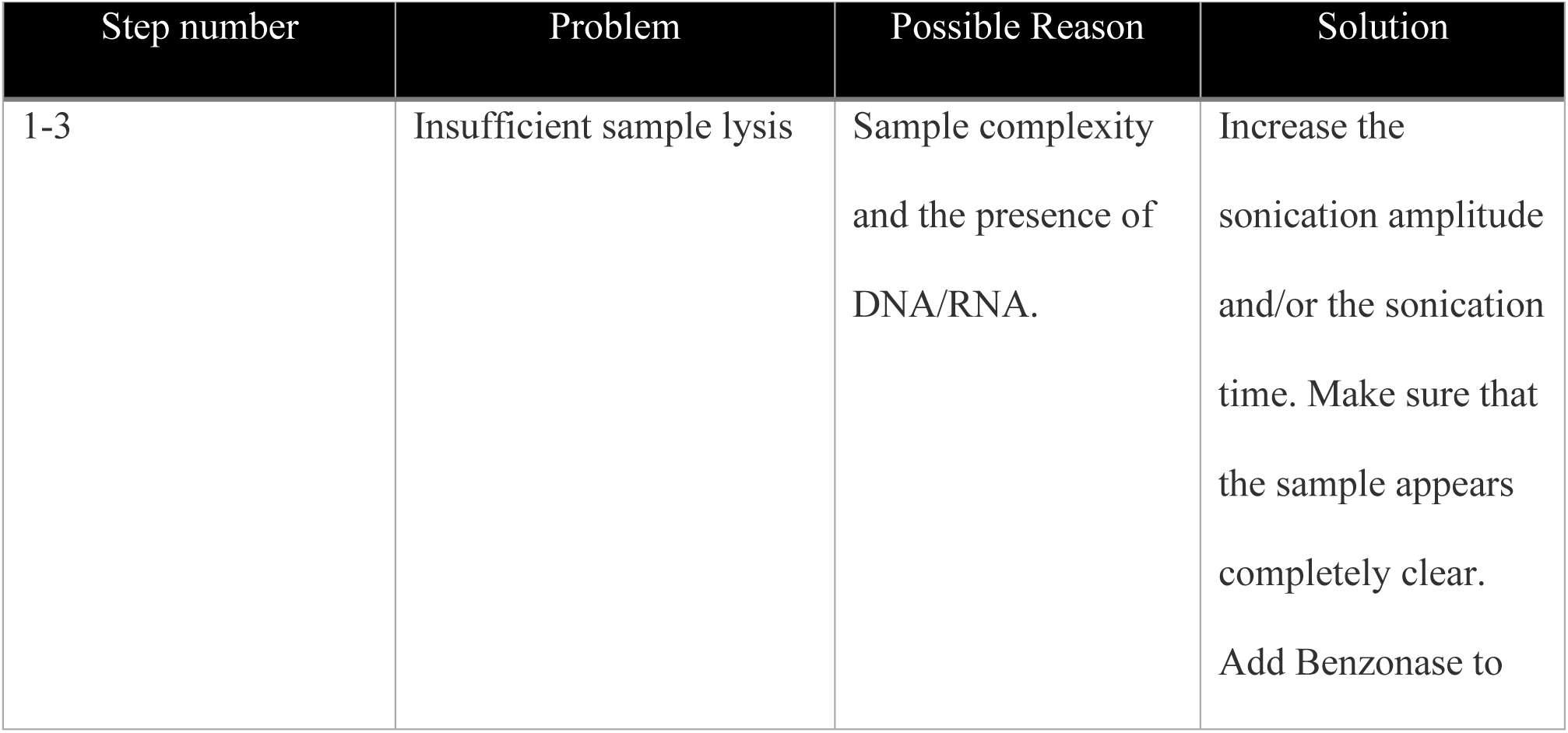
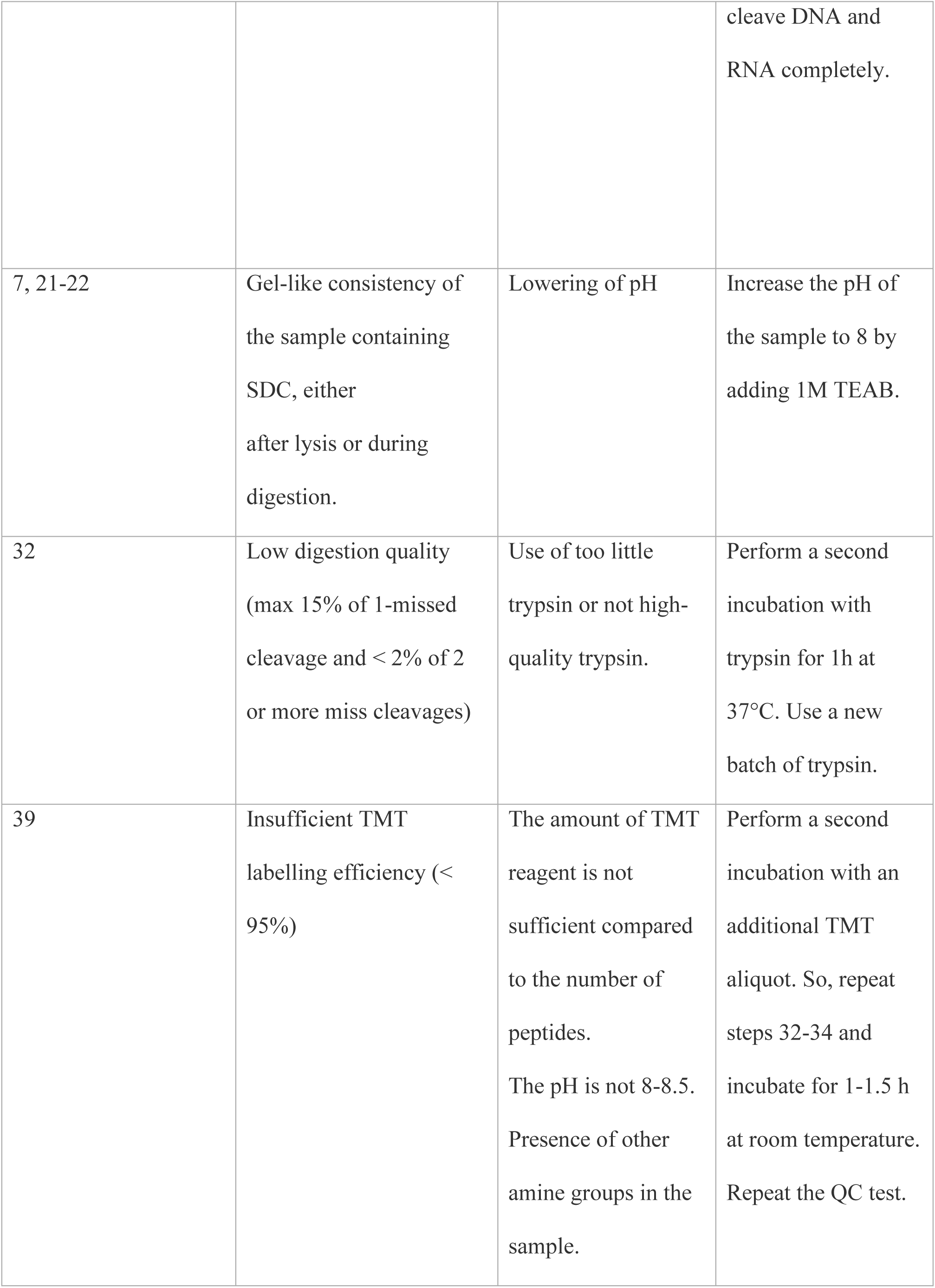
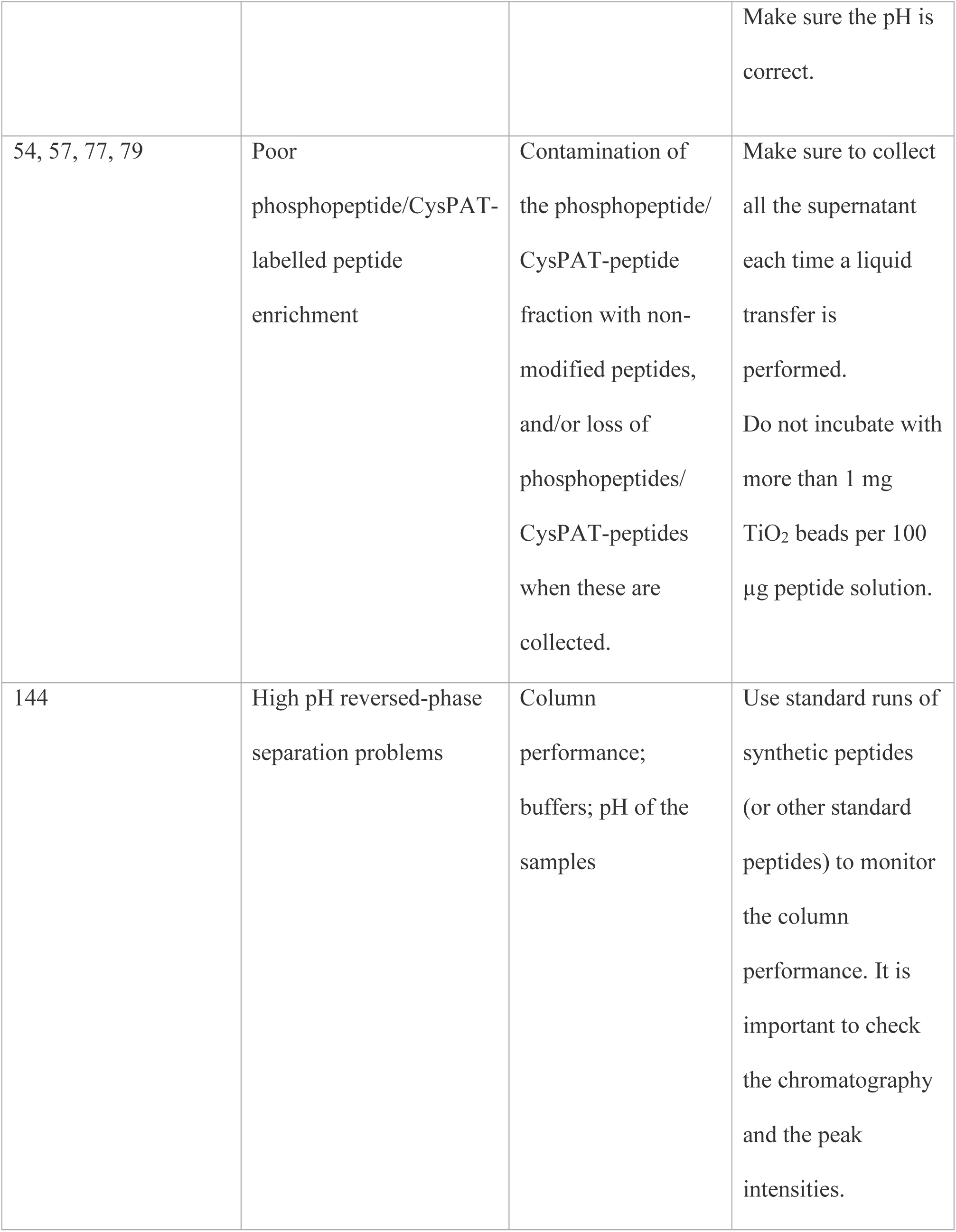
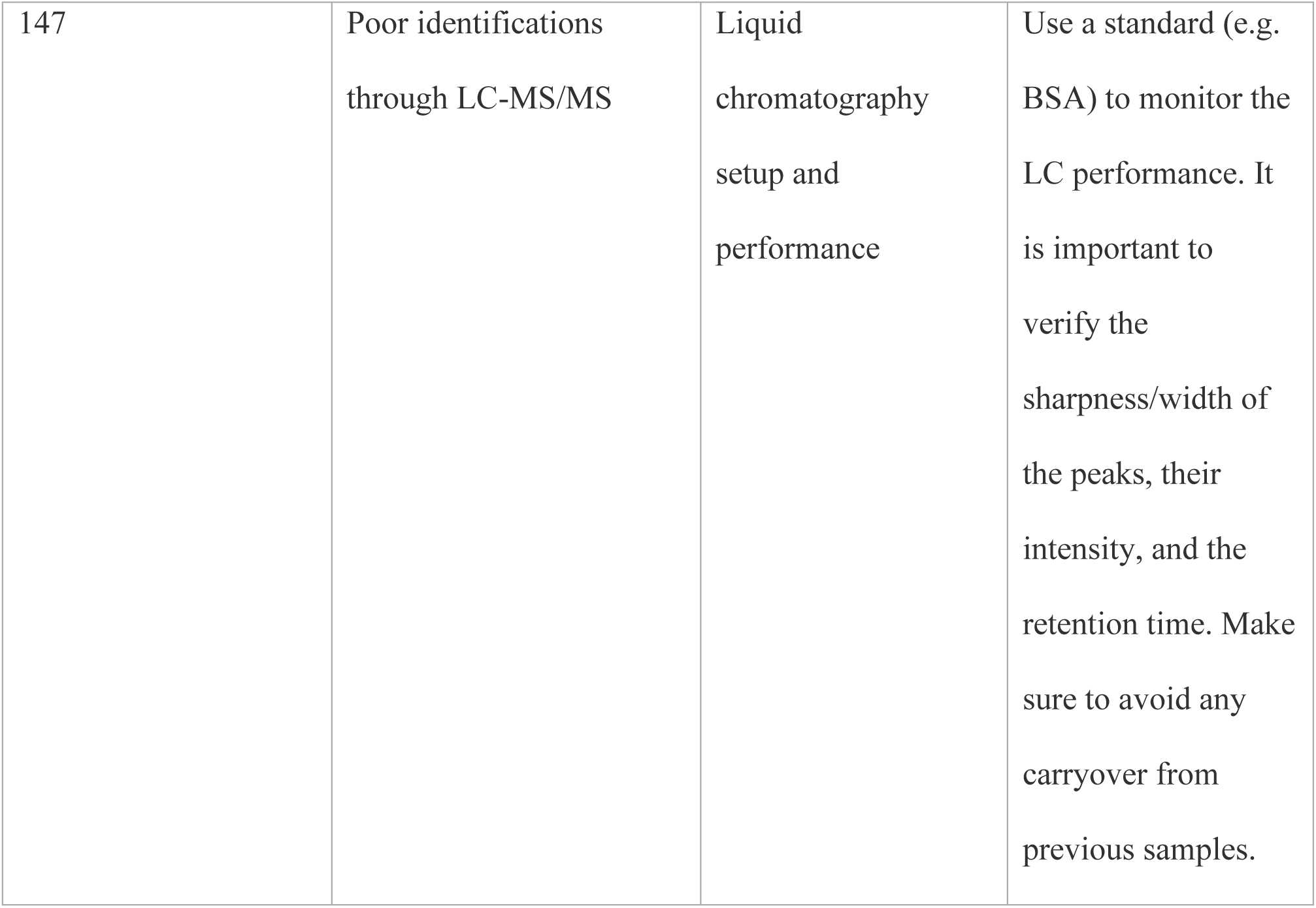

#### Anticipated results

The method described here (illustrated in **Figure 1**) enables the simultaneous and global profiling of the proteome and several PTMs from cell or tissue samples. To provide a demonstration of its performance, the protocol was applied on unguided neural organoids [66] at various time points (day 130, 140, 150, and 160). From a single TMT16 plex experiment and using 1.8 mg as total starting material (150 µg protein per condition, using 12 out of 16 TMT channels), we quantified 10,413 proteins of which 7,797 were identified in the non-modified dataset with at least 2 unique peptides per protein (**Figure 2a**). In the PTM-enriched samples, we quantified a total of 19,655 phosphopeptides from 4,199 phosphoproteins, 1,716 glycopeptides from 797 formerly N-linked sialylated glycoproteins, 9,723 peptides from 4,702 free cysteine-containing proteins, 28,876 peptides from 8,203 reversibly modified cysteine-containing proteins, and 771 peptides from 409 lysine acetylated proteins (**Figure 2a+b**)(**Supplementary Tables CBO study**). Metabolomics analysis additionally resulted in the identification and quantification of 4,493 metabolites of which 2,969 metabolites were quantified in all samples. A total of 331 of the 2,969 metabolites present in all neural organoid samples could be annotated to Metabolomics Standard Initiative (MSI) level 3 [67] (**Figure 2c**). A large proportion of the identified proteins were found in one or more of the PTM datasets (**Figure 2a + Supplementary Figure S1**), while some proteins were exclusively identified in a single PTM dataset (**Supplementary Figure S1**). This illustrates that proteins exist in multiple proteoforms [1] and that enrichment of multiple PTMs from a biological sample increases the global coverage of the proteome. In addition, we found that our approach could identify 69% of brain-specific proteins (**Figure 2d**), including known neuronal and astrocytic markers such as doublecortin (DCX), microtubule-associated protein 2 (MAP2), synaptophysin (SYP), and glial fibrillary acidic protein (GFAP). Thus, the protocol allows for the identification and quantification of not only a substantial proportion of the proteome and PTMs but also covers proteins that are biologically relevant in the context of tissue structure and function.

**Figure 2.**
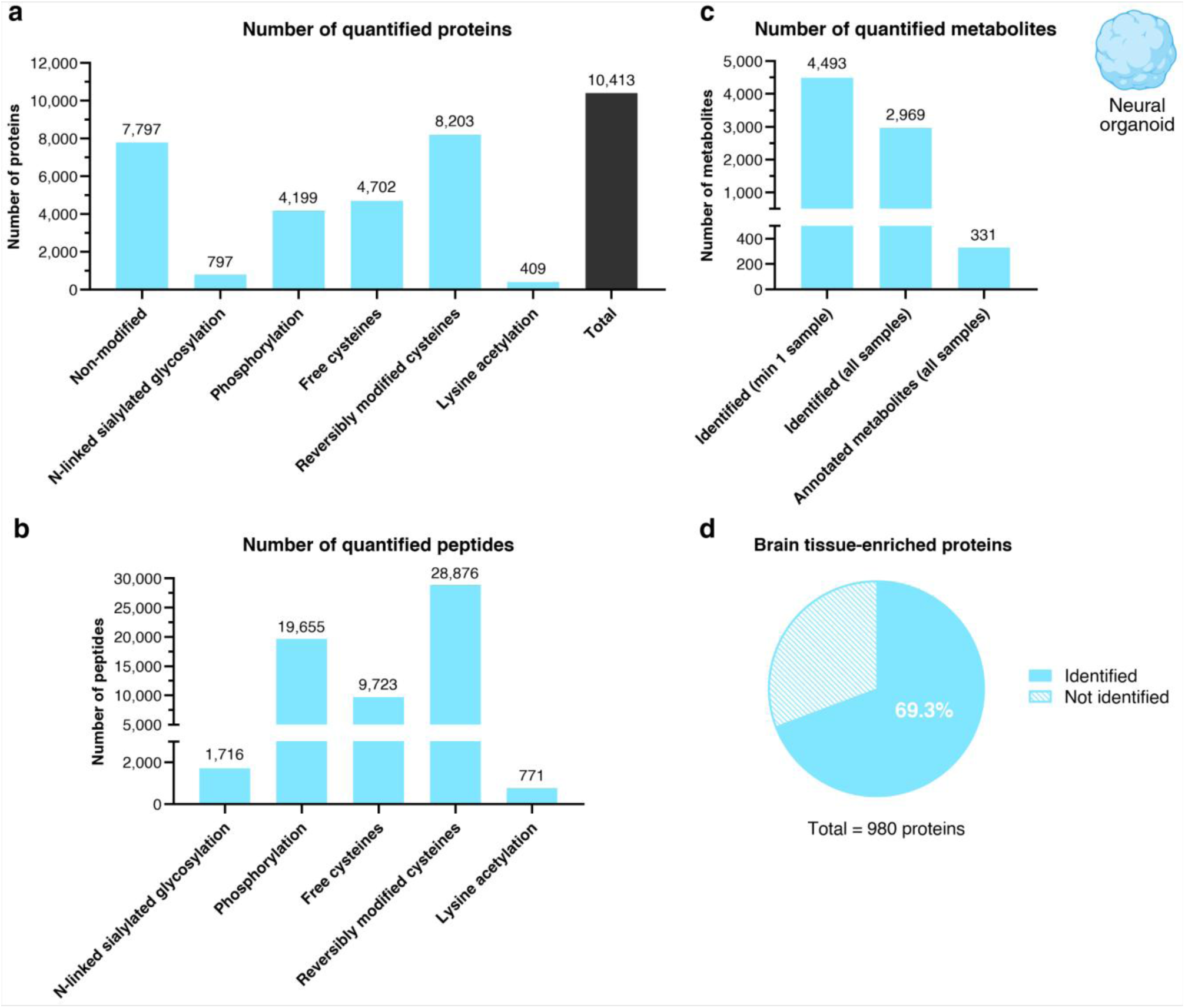
| Coverage of protein, post-translational modification (PTM) and metabolite abundances in unguided neural organoids. **A-c**, Bar charts illustrate the number of quantified non-modified and PTM proteins (**a**), the number of quantified PTM peptides (**b**), and the number of quantified metabolites – measured with the *Metabolomics setup* described in the sections *Mass spectrometer methods* and *Database searching* (**c**). Blue bars indicate the number of identified unique proteins (**a**), peptides (**b**), and metabolites (**c**) in each dataset, while the dark grey bar in (**a**) indicate the total number of proteins identified across the proteomic and PTMomic datasets. **D**, the proportion of brain-specific proteins identified in unguided neural organoids across the proteomic and PTMomic datasets.

To provide additional examples of the performance of this method on other tissue types, the protocol was applied on mouse heart and liver samples. In both experiments, a TMT16 plex analysis was performed, using 100 µg of proteins for each sample/channel as starting material. To demonstrate that the protocol is easily adaptable to the needs of different experiments, such as adding and/or skipping one or more PTM enrichment steps, we added an extra step of enrichment of S-palmitoylated peptides (as described in [52]) for the mouse heart tissue (adding to a total of 7 PTMs analyzed), whereas for the mouse liver tissue, we skipped the lysine acetylation enrichment step (corresponding to analysis of a total of 5 PTMs). As shown in **Figure 3**, the protocol gave comparable results with the neural organoid experiment and a high coverage of both the proteome and PTMome. In total, 7,492 and 7,828 proteins were quantified across the proteomic and PTMomic datasets in the mouse liver and heart tissue, respectively (**Figure 3a+c**). In terms of PTM proteins, we identified and quantified 3,097 and 2,327 phosphorylated proteins, 4,740 and 5,671 reversibly modified cysteine-containing proteins, 4,939 and 5,238 free cysteine-containing proteins, as well as 778 and 796 formerly N-linked sialylated glycoproteins, respectively, in mouse liver and heart (**Figure 3a+c**). In addition, 185 lysine acetylated proteins and 3,595 depalmitoylated proteins were quantified in mouse heart tissue (**Figure 3c**) (see lists in **Supplementary Tables Heart mouse study and Supplementary Tables Liver mouse study)**. As for the neural organoid samples, most proteins identified in mouse liver and heart samples carried PTMs, and some proteins were uniquely identified only in PTM-enriched datasets. Only a minority of the proteins identified for each tissue were identified across all datasets (**Supplementary figure S2+S3**). The subcellular localization of the identified proteins, including nuclear, cytosolic, mitochondrial, secreted, and membrane-associated -was further evaluated for each PTM, illustrating how different cellular compartments are proportionally represented across protein modifications (**Supplementary figure S4**). For both liver and heart tissue samples, around 70% of tissue-specific proteins were identified and quantified. For liver tissue, this included proteins important for liver physiological functions such as plasma proteins (APOB, C2, FGA), metabolic enzymes (HAO1, RDH16, ALDOB), proteins involved in bile synthesis (SLC27A5, BAAT), and transporter proteins (ABCB11, SLC2A2), whereas for heart tissue it included proteins related to muscular contraction (MYH7, ACTC1, TNNI3), electrolyte homeostasis (NPPA, CASQ2, PLN), and intercalated discs (ATP1A3, CDH2).

**Figure 3.**
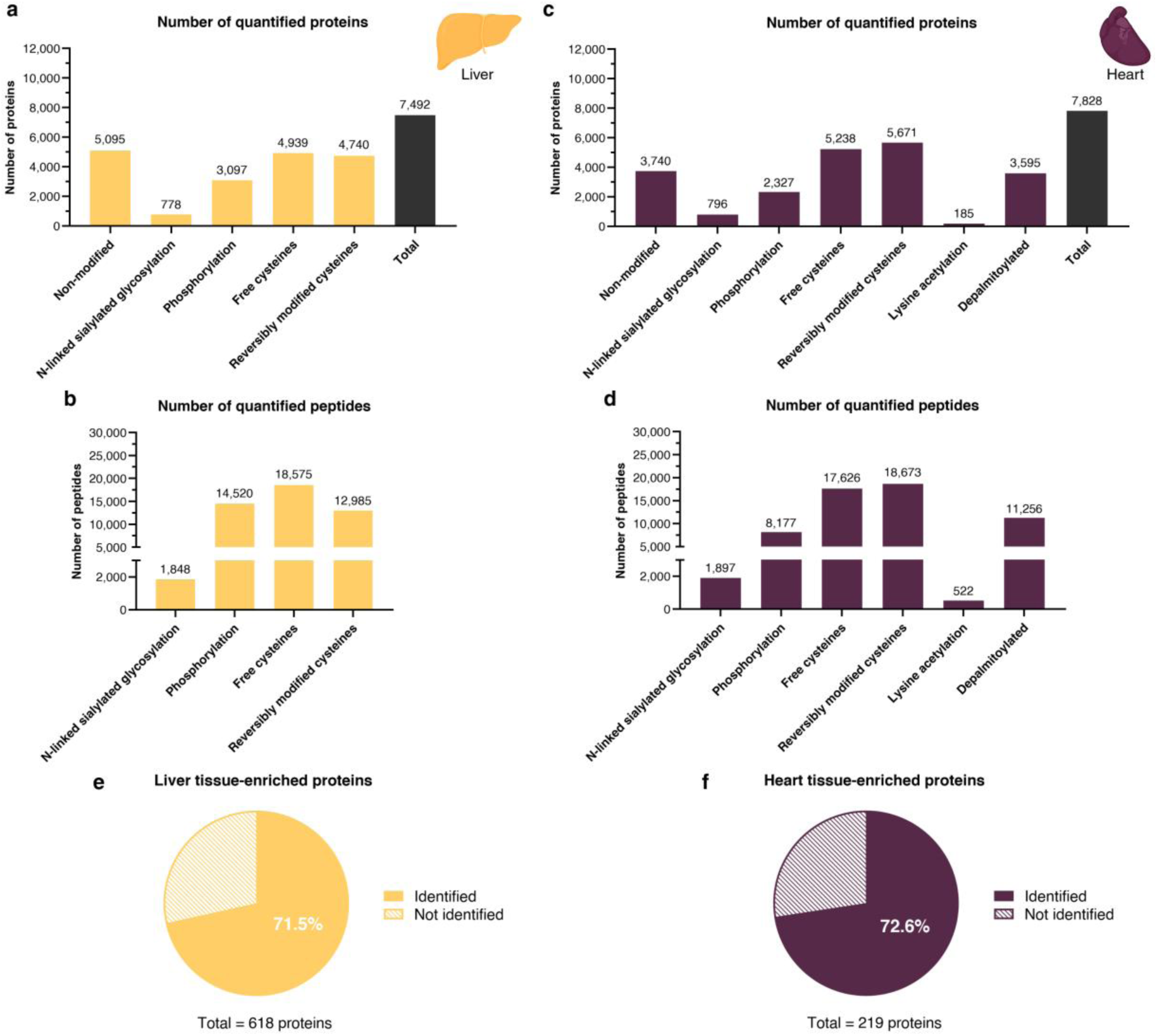
| Coverage of protein and post-translational modification (PTM) abundances in liver and heart tissue samples. **a-d**, Bar charts illustrate the number of quantified non-modified and PTM proteins (**a+b**) and the number of quantified PTM peptides (**c+d**) in liver and heart tissue. Yellow bars indicate the number of identified unique proteins (**a**) and peptides (**c**) in each dataset for liver tissue, while purple bars indicate the number of identified unique proteins (**b**) and peptides (**d**) in each dataset for heart tissue. Dark grey bars in **a** and **b** indicate the total number of proteins identified across the proteomic and PTMomic datasets. **e-f**, The proportion of liver-specific proteins (**e**) and heart-specific proteins (**f**) identified in liver and heart tissue, respectively, across the proteomic and PTMomic datasets.

## Acknowledgements

The present study was supported by the Lundbeck foundation (R336-2020-1113) and the Villum Center for Bioanalytical Sciences at SDU. We also thank former member of the Larsen group at the Department of Biochemistry and Molecular Biology at the University of Southern Denmark for various contributions to development of methods for PTMs over the last 20 years.

## Author contributions

**Lucrezia Criscuolo:** Formal analysis, Investigation, Data curation, Writing – original draft, Writing – review and editing, Visualization. **Sofie B. Elmkvist**: Data curation, Data analysis, Writing – original draft, Visualization. **Arkadiusz Nawrocki**: Mouse tissue analysis, Data curation, Methodology. **Lene A. Jakobsen**: Mouse tissue analysis, Methodology. **Pia Jensen**: Data curation, Methodology, Writing – review and editing. **Peter T. Jensen**; Methodology. **Honggang Huang**: Methodology. **Jesper F. Havelund**: Methodology. **Nils J. Færgeman**: Methodology, Writing – review and editing. **Giuseppe Palmisano:** Methodology, Resources, Writing – review and editing. Helle Bogetofte: Methodology, Writing – review and editing. **Martin R. Larsen:** Conceptualization, Methodology, Resources, Writing – review and editing, Supervision, Project administration, Funding acquisition.

---

**Figure S1.**
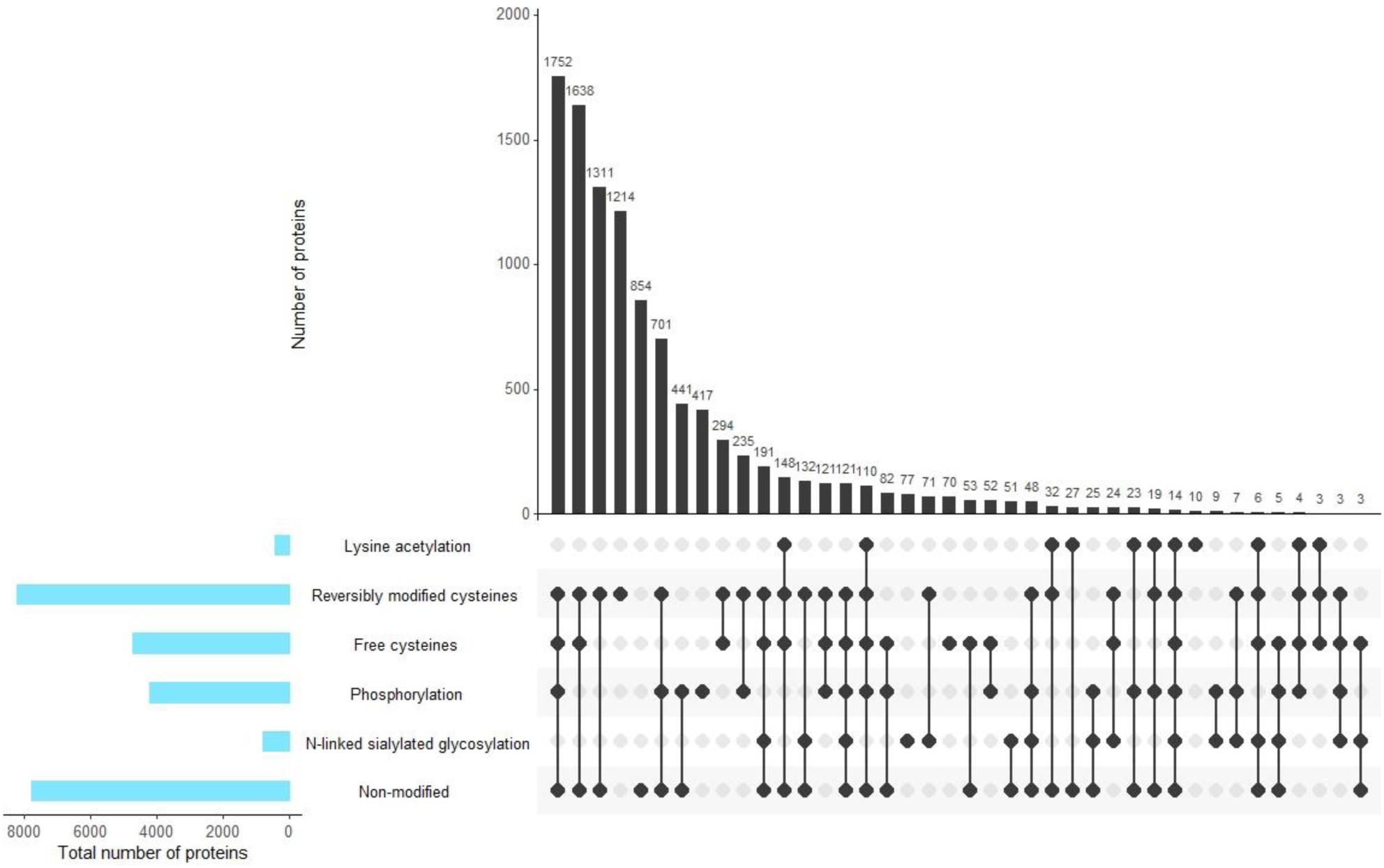
| Overlap in proteins identified in proteomics and PTMomics data in cerebral organoids.

**Figure S2.**
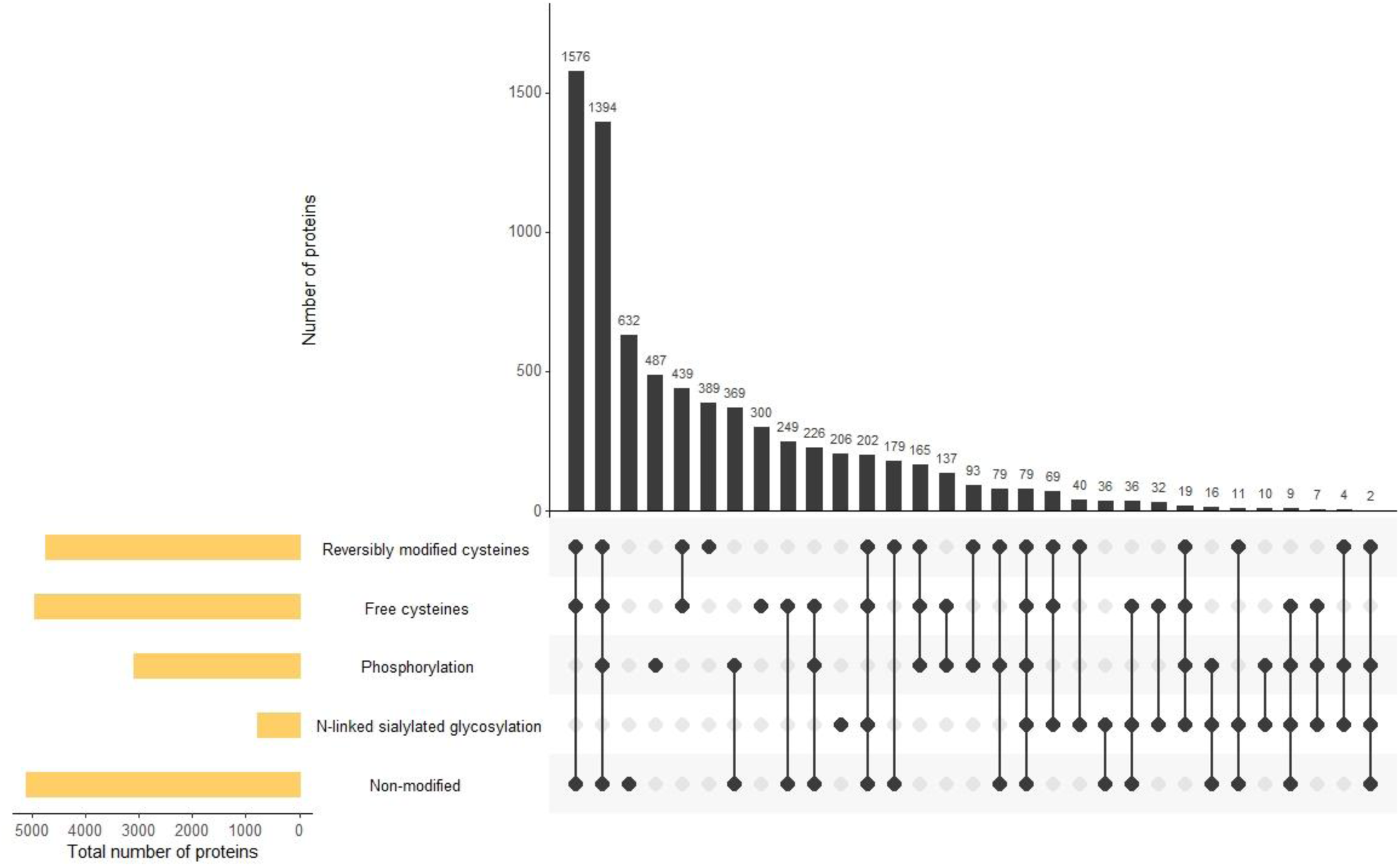
| Overlap in proteins identified in proteomics and PTMomics data in liver tissue.

**Figure S3.**
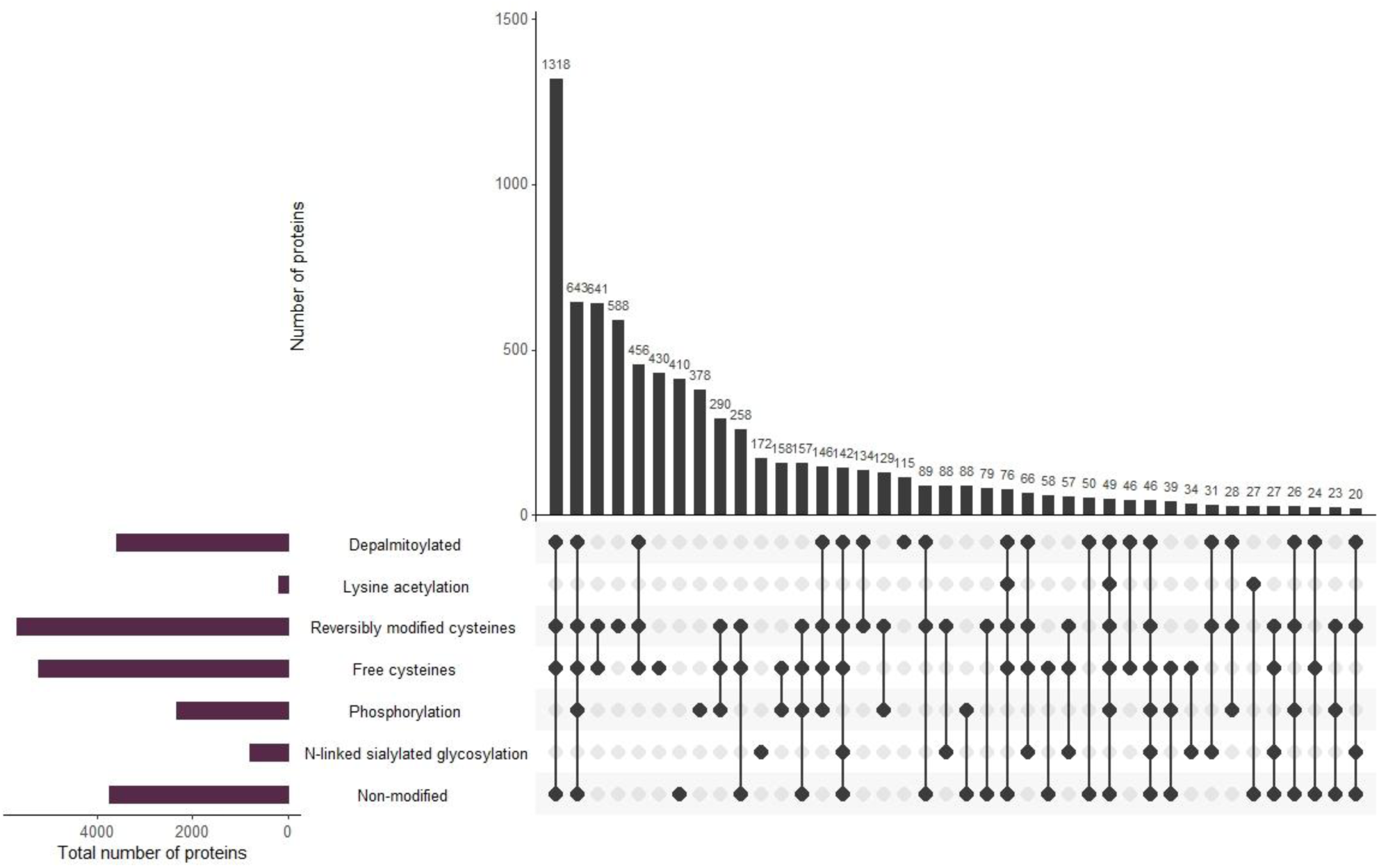
| Overlap in proteins identified in proteomics and PTMomics data in heart tissue.

**Figure S4.**
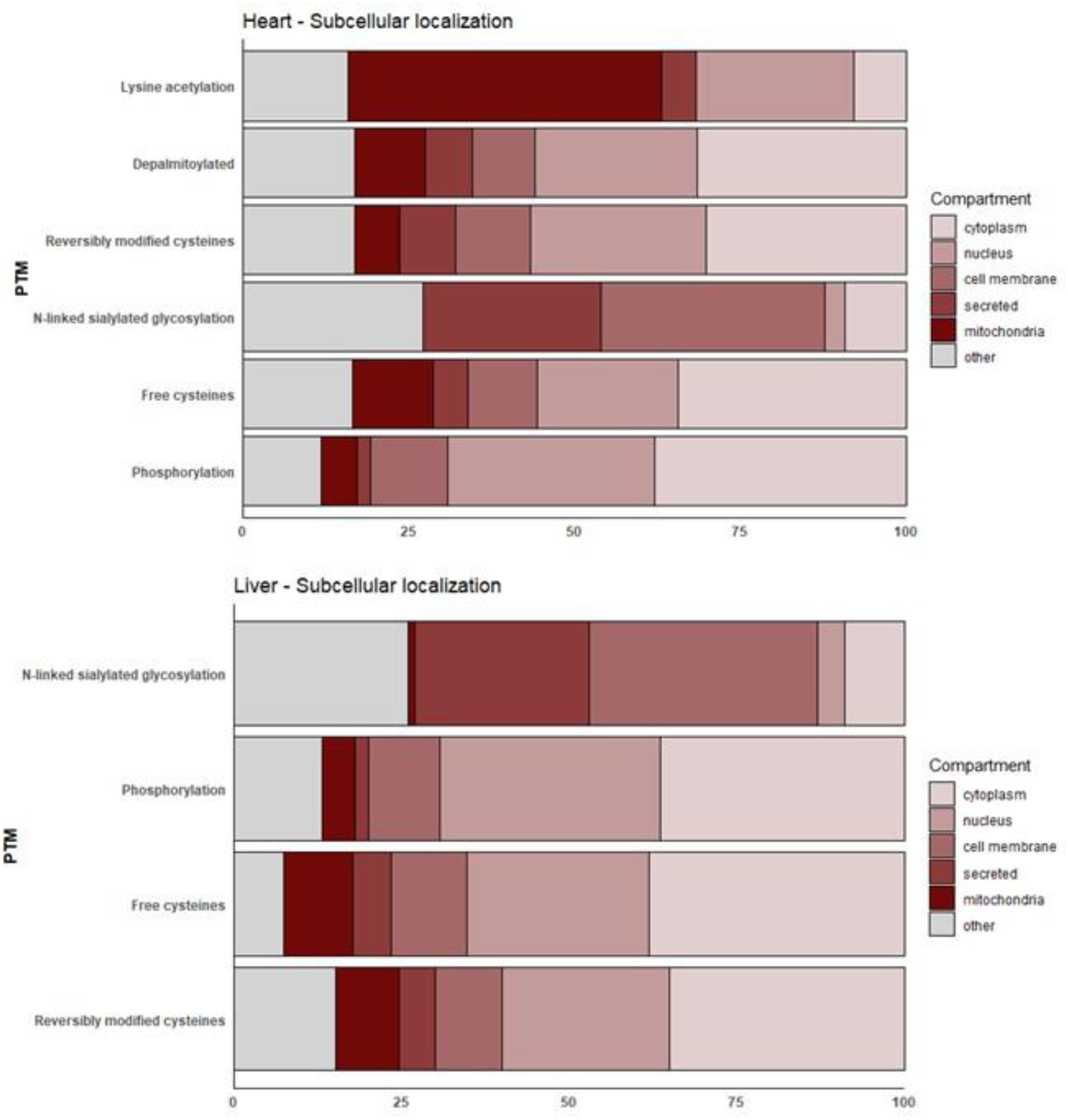
| Subcellular localization of proteins identified in PTMomics data in heart and liver tissue. Localization was assigned based on UniProt annotations, which were grouped into nucleus, cytoplasm, mitochondria, secreted, membrane-associated proteins, and others. Data are shown as percentages for each PTM fraction.

